# Aperiodic neural activity links electromagnetic and hemodynamic representations of domain-general cognitive demand across the cortical hierarchy

**DOI:** 10.64898/2026.08.03.742517

**Authors:** Runhao Lu, Moataz Assem, Xiaobo Liu, John Duncan, Alexandra Woolgar

## Abstract

The human brain demonstrates remarkable flexibility and capacity for domain-general cognitive control, allowing us to perform diverse and complex tasks. Central to this ability is the multiple-demand (MD) network, a domain-general system that is robustly engaged during demanding tasks in fMRI studies. However, the electrophysiological signatures underlying these domain-general responses remain elusive. While recent research has implicated aperiodic neural activity as a promising candidate, the limited spatial resolution of non-invasive electrophysiology has left it unresolved how this aperiodic signal relates to demand-related activity within the MD network and whether this relationship reflects a broader organizational principle across the cortex. To address these questions, we used a multimodal fusion framework to integrate fMRI and magnetoencephalography (MEG) data acquired while participants performed a diverse set of cognitive control tasks. We found that raw MEG-fMRI correspondence was strongest in unimodal sensorimotor cortices and progressively decreased toward transmodal association cortex during cognitive control tasks, revealing a hierarchical decline in correspondence between the electromagnetic and hemodynamic signals measured by these technologies. However, the proportion of this variance that was attributable to cognitive demand and carried by aperiodic signals showed the reverse gradient, systematically increasing along the sensorimotor-association axis. In particular, in the MD network, aperiodic broadband power showed the strongest demand-specific cross-modal commonality, outperforming canonical oscillatory components. These findings reveal two opposing hierarchical gradients: overall MEG–fMRI correspondence across all electrophysiological signals decreased toward association cortex, whereas the proportion attributable to aperiodic signals associated with cognitive demand increased. Our results identify aperiodic neural activity as a key electrophysiological substrate of cognitive control and a bridge linking electromagnetic and hemodynamic representations across the cortical hierarchy.

## Introduction

Adaptive human behaviour relies on domain-general cognitive systems that flexibly support diverse tasks through shared neural mechanisms. A substantial body of research has linked this domain-general capacity to a specific neural system commonly referred to as the multiple-demand (MD) brain network (Assem, Glasser, Van Essen, & Duncan, 2020; Duncan, Assem, & Shashidhara, 2020; Fedorenko, Duncan, & Kanwisher, 2013; Schultz, Ito, & Cole, 2022), and also known as the frontoparietal control network (Dixon et al., 2018; Murphy, Bertolero, Papadopoulos, Lydon-Staley, & Bassett, 2020; Nee, 2021; Yeo et al., 2011), task-positive network (Fox et al., 2005; Spreng, Wojtowicz, & Grady, 2010), or cognitive control network (Cole & Schneider, 2007; Gratton, Sun, & Petersen, 2018). This network is distributed across widespread brain regions, particularly within the frontal and parietal cortices. In fMRI studies, it is characterized by its robust activation across diverse cognitive tasks when contrasting hard versus easy conditions (Assem, Shashidhara, Glasser, & Duncan, 2024; Fedorenko et al., 2013) and flexible multivariate representation of a wide range of task features (e.g., Zheng, Lu, and Woolgar (2024)). However, the underlying electrophysiological signatures that bridge these hemodynamic responses to direct neurophysiological activity remain elusive.

Although many electrophysiological studies have traditionally investigated how periodic neural oscillations in specific frequency bands respond to cognitive demand and potentially relate to frontoparietal MD network (Chikhi, Matton, & Blanchet, 2022; Pavlov & Kotchoubey, 2022), the results remain largely inconclusive. Findings across canonical frequency bands often yield contradictory directions of change, and their spatial localizations frequently diverge from the widespread activation pattern characteristic of the MD network (Lu, 2025). Beyond oscillatory responses, recent electrophysiological findings suggest that the aperiodic (1/f-like) neural activity may reflect widespread brain network excitability and the balance of synaptic excitation and inhibition (E/I balance) (Cohen Kadosh, 2025; Gao, Peterson, & Voytek, 2017; Waschke et al., 2021), potentially bridging electrophysiological signals and the network-level activations detected by fMRI (Jacob, Roach, Sargent, Mathalon, & Ford, 2021; Lu, 2025; Wen & Liu, 2016a). Notably, emerging evidence suggests that aperiodic activity plays important roles in a wide range of cognitive processes, such as selective attention (Lu, Pollitt, & Woolgar, 2025), working memory (Bender, Zhao, Vogel, Awh, & Voytek, 2025; Donoghue et al., 2020; van Engen et al., 2026), inhibitory control (Jia et al., 2024), and task switching (Yan et al., 2024). In a recent study, we examined the roles of oscillatory and aperiodic neural activity in coding task demand across six diverse cognitive tasks (Lu, Dermody, Duncan, & Woolgar, 2024), and found that aperiodic broadband power (BB) was the most robust index for decoding cognitive demand that generalizes across distinct task domains. These results indicate that aperiodic activity potentially serves as a fundamental electrophysiological substrate for flexible, domain-general cognitive control observed in the human brain.

Despite these parallel lines of evidence from fMRI and electrophysiology, how these two imaging modalities spatially correspond, and specifically how their relationship is organized across the cortical hierarchy during cognitive control, remains unknown. Recent macroscale mapping studies have established a principal gradient of cortical organization, spanning from unimodal sensorimotor areas to transmodal association cortices (Bernhardt, Smallwood, Keilholz, & Margulies, 2022; Margulies et al., 2016; Sydnor et al., 2021), in which the MD network is prominently distributed across these higher-order transmodal regions. Along this sensorimotor-association (S-A) axis, recent studies observed a macroscale pattern of increased decoupling between electrophysiological and haemodynamic responses. While unimodal cortices exhibit strong coupling between structural and functional networks (Baum et al., 2020; Vazquez-Rodriguez et al., 2019) as well as between electromagnetic and hemodynamic activities (Shafiei, Baillet, & Misic, 2022), this correspondence profoundly decouples in transmodal association cortices. However, this hierarchical decoupling has been largely established during resting-state, leaving it unclear whether it is maintained during active task states such as when people engage in cognitive control. Moreover, if the resting state data reflect a “baseline” decoupling motif in transmodal regions, identifying the electrophysiological basis of cognitive control will require a framework that accounts for both global cortical hierarchies and local network specificity. While our previous magnetoencephalography (MEG) findings highlighted the importance of BB activity in domain-general cognitive control, the limited spatial resolution of MEG made it difficult to determine whether this BB-mediated response is locally confined to the MD network or reflects a broader, hierarchical property across the association cortex.

To address these challenges, we employed a multimodal fusion framework (e.g., Cichy & Oliva, 2020; Karimi-Rouzbahani, Rich, & Woolgar, 2026; Moerel, Rich, & Woolgar, 2024) to integrate MEG and fMRI data acquired while participants performed the same set of cognitive tasks with varying levels of demand. This allowed us to ask three primary questions: First, does the macroscale decoupling between electromagnetic and hemodynamic signals persist during active cognitive control tasks? Second, which component of the electrophysiological signal from MEG best captures demand-related variance that is shared with BOLD responses from the MD network? Third, to what extent does this relationship reflect a broader organizational pattern across the cortical hierarchy?

By mapping cross-modal correspondence at two levels: model-free raw representational similarity and model-based demand-specific commonality, we identified two opposing hierarchical gradients. Addressing our first question, the model-free analysis revealed that raw MEG-fMRI correspondence systematically decreased along the S-A axis across all electrophysiological signals, mirroring resting-state decoupling (Baum et al., 2020; Shafiei et al., 2022; Vazquez-Rodriguez et al., 2019). For the second question, we found that within the MD network, aperiodic BB activity was the most robust predictor of demand-specific cross-modal commonality, significantly outperforming canonical oscillations, thereby positioning aperiodic dynamics as a key electrophysiological substrate for domain-general cognitive control. Finally, addressing our third question, our whole-cortex analysis revealed that this task-driven functional convergence is not exclusively confined to the MD network. Instead, isolating representational variance specifically related to cognitive demand revealed an opposing hierarchical gradient uniquely for aperiodic signals: the proportion of cross-modal shared variance attributable to cognitive demand systematically increased along the S-A axis, peaking in the association cortex. Together, these findings demonstrate that while electromagnetic and hemodynamic signals in higher-order regions are intrinsically decoupled, they exhibit a proportional representational specialization for task demands, with aperiodic activity serving as the essential bridge across the cortical hierarchy.

## Results

To investigate domain-general neural responses to diverse cognitive demand, we employed three cognitive tasks in both fMRI and MEG studies. Owing to practical constraints in participant recruitment and longitudinal follow-up, the two modalities were acquired in independent samples, with 36 adults (age range: 21-49 years; 20 females, 16 males) in the fMRI study and 43 adults (age 18-39 years, 31 females and 12 males) in the MEG study. To mitigate the inherent limitations of comparing independent samples, we utilized a robust statistical framework for the fusion analyses, incorporating a bootstrapping procedure to derive stable group-level representational estimates and permutation testing to assess the significance of cross-modal correspondence.

Across both modalities, participants performed a working memory task (WM), a switching task (SWIT), and a multi-source interference task (MSIT) (Figure 1A). Following a 3*2*2 design, each task involved two types of stimulus content (alphanumeric or colour stimuli) and two levels of demand (hard vs. easy). For the WM task, demand was manipulated via memory load (4 vs. 2 items). In the SWIT task, demand was defined by trial type, contrasting switch trials (high demand) with repeat trials (low demand). For the MSIT task, demand was manipulated by spatial congruency and distractor interference, with incongruent trials serving as the high-demand condition. Detailed results of the MEG study have been reported in Lu et al. (2024).

**Figure 1.**
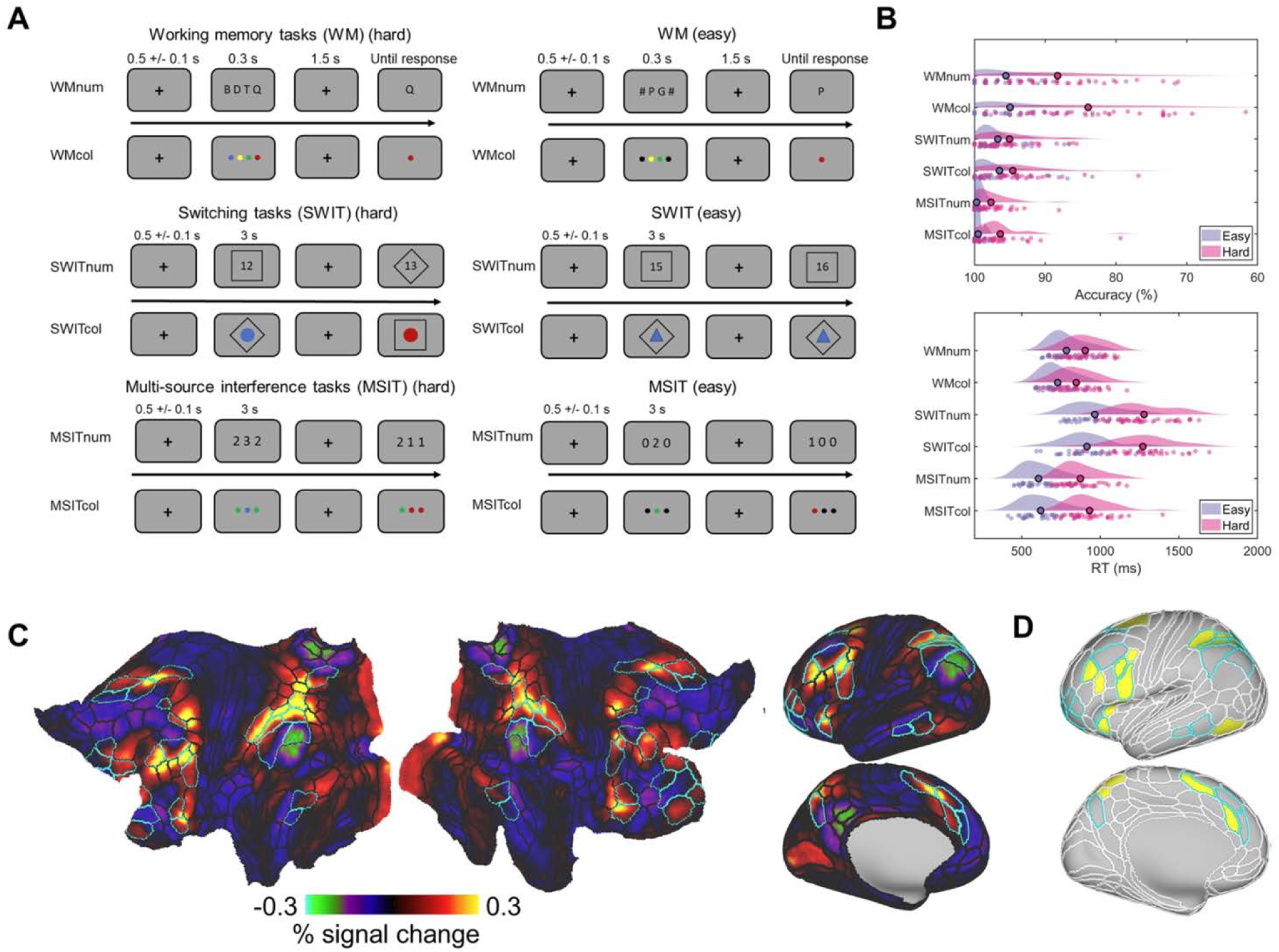
Experimental paradigm, behavioural and activation results of the fMRI study. (A) Experimental design of the working memory task (WM), the switching task (SWIT), and the multi-source interference task (MSIT). Each task had two versions with either alphanumeric or colour stimuli as task content. (B) Behavioural accuracy (top) and response time (RT; bottom) in easy and hard conditions for each subtask. Each individual dot in the raincloud plots represents a participant and the bolded dot shows the mean. Repeated measures ANOVAs (task demand * task content) showed that the main effects of task demand (hard vs. easy) were significant for both accuracy and RT for all 6 subtasks (all *p*s < 0.001). WM: Working memory task; SWIT: Switching task; MSIT: Multi-source interference task; num: Alphanumeric task; col: Colour task. (C) Group average univariate activation maps on flattened (left) and inflated cortical surfaces (right) for the hard > easy contrast, averaged across six conditions [3 tasks (WM, SWIT, and MSIT) × 2 contents (alphanumeric and colour)]. (D) Empirical MD patches (yellow) identified using the current dataset. The cortical surface was parcellated into 360 regions (180 per hemisphere) based on the HCP-MMP1.0 Parcellation (outlined in white) (Glasser et al., 2016). Cyan borders surround the extended MD areas defined in (Assem et al., 2020). Only one hemisphere is shown for the inflated surfaces in (C) and (D), given that the activation patterns were highly consistent across hemispheres and the empirical MD network was defined bilaterally.

### Behavioural results

To verify the demand manipulation in the fMRI study, we conducted 2 (task demand: hard vs. easy) × 2 (task content: alphanumeric vs. colour) repeated measures ANOVAs on behavioural accuracy and reaction time (RT) for each task. As shown in Figure 1B, we found a robust main effect of task demand in all three tasks for both accuracy and RT (all *F*s > 19.84, *p*s < 0.001, *η*_p_^2^ > 0.36; see Table S1 for descriptive statistics). Participants were consistently more accurate and faster in the easy compared to the hard conditions, confirming the efficacy of our demand manipulation across diverse task domains.

We further examined the effects of task content and its interaction with demand for each task separately. For the WM task, we found a significant main effect of task content and a significant interaction between demand and content for behavioural accuracy [content effect: *F* (1,35) = 6.39, *p* = 0.02, *η*_p_^2^ = 0.15; interaction: *F* (1,35) = 5.72, *p* = 0.02, *η*_p_^2^ = 0.14]. Participants were significantly less accurate in the colour WM than in the alphanumeric WM task in the hard condition [*F* (1,35) = 8.40, *p* = 0.006] but not in the easy condition [*F* (1,35) = 0.38, *p* = 0.54], though the difficulty effects remained significant for both modalities (both *F*s > 42.51, *p*s < 0.001). For RT, we found a significant content effect [*F* (1,35) = 41.94, *p* < 0.001, *η*_p_^2^ = 0.55]. Participants had faster RT in the colour compared to alphanumeric WM task, which may have reflected a speed-accuracy trade-off. No interaction effects were found for RT [*F* (1,35) = 0.001, *p* = 0.99, *η*_p_^2^ < 0.001].

For the SWIT task, we did not find a significant content effect or interaction between demand and content for behavioural accuracy [content effect: *F* (1,35) = 0.29, *p* = 0.60; interaction: *F* (1,35) = 0.14, *p* = 0.71]. However, for RT, we found a significant interaction effect [*F* (1,35) = 9.84, *p* = 0.003, *η*_p_^2^ = 0.22]. Participants had slower RT in the alphanumeric compared to colour SWIT tasks only in the easy condition [*F* (1,35) = 7.78, *p* = 0.01]. The difficulty effects were significant for both modalities (Fs > 652.02, ps < 0.001).

For the MSIT task, there was a significant main effect of task content and a marginally significant interaction between demand and content for accuracy [content effect: *F* (1,35) = 7.77, *p* = 0.01, *η*_p_^2^ = 0.18; interaction: *F* (1,35) = 4.05, *p* = 0.05, *η*_p_^2^ = 0.10]. This interaction indicated that the performance drop from the easy to the hard condition was slightly larger for the colour MSIT compared to the alphanumeric MSIT. Although our results also showed that participants were significantly less accurate in the colour MSIT than in the alphanumeric MSIT task across conditions [easy condition: *F* (1,35) = 6.55, *p* = 0.02; hard condition: *F* (1,35) = 5.94, *p* = 0.02], these were very small differences as accuracy was near ceiling (e.g., mean accuracy of easy alphanumeric and colour task was 99.72% and 99.49%, respectively; also see Table S1). Again, the demand effects were significant for both the alphanumeric and colour MSIT tasks (both *F*s > 25.17, *p*s < 0.001). For RT, we found a significant interaction effect [*F* (1,35) = 40.56, *p* < 0.001, *η*_p_^2^ =0.54]. Participants had slower RT in the colour compared to alphanumeric MSIT tasks only in the hard condition [*F* (1,35) = 24.68, *p* < 0.001]. The difficulty effects were again significant for both modalities (*F*s > 652.02, *p*s < 0.001).

In summary, the behavioural results largely replicated results from the MEG study (Lu et al., 2024) and confirmed that hard conditions were more cognitively demanding than easy conditions across all tasks, as expected.

### Domain-general task demand activation and empirical MD network definition

To identify brain regions sensitive to cognitive demand, we first examined the univariate fMRI activation for the hard > easy contrast, averaged across all six subtasks (Figure 1C; see Figure S1 for thresholded map with Bonferroni correction). Consistent with the well-established MD network (Assem et al., 2020; Duncan et al., 2020), increased task demand robustly recruited a distributed set of frontal and parietal regions, including the dorsal and mid-frontal cortex, anterior insula, intraparietal sulcus, and dorsomedial parietal regions. This activation pattern was highly consistent across the three task domains (WM, SWIT, and MSIT) and both stimulus contents (alphanumeric and colour), confirming the domain-general nature of these responses (Figure S2).

While the spatial distribution of demand-related activation broadly replicated the canonical MD patches defined by (Assem et al., 2020), we observed some shifts in the exact peak locations, particularly within the parietal and posterior temporal cortices. For example, we observed more dorsal but less ventral activation around the intraparietal sulcus, the posterior temporal patch in our results appeared more posterior, and the anterior frontal patch appeared to show weaker activation, particularly in the MSIT tasks. These shifts could reflect the different participants and/or tasks used in this study. To accommodate these spatial characteristics, we defined an empirical MD network as a single, combined region of interest (ROI) to optimize the spatial precision of our subsequent model-based fusion analyses. To do this, we identified cortical areas showing significant activation (*p* < 0.05, Bonferroni corrected) in at least five of the six subtask contrasts (see Methods for details). This procedure yielded a robust set of 22 bilateral areas (Figure 1D and Figure S3) that were united to form a single MD ROI for subsequent analysis.

### RDM construction and split-half reliability

To characterize neural representations across imaging modalities, we employed a representational similarity analysis (RSA) framework. We first constructed representational dissimilarity matrices (RDMs) for each modality based on 12 experimental conditions (3 tasks × 2 task demands × 2 task contents) (Figure 2A, B). For the MEG data, we removed event-related fields from each trial to focus on induced responses and then utilized irregular resampling auto-spectral analysis (IRASA; (Wen & Liu, 2016b)) to separate the neural signal into specific oscillatory (theta, alpha, and beta) and aperiodic (BB, slope, and intercept) components (Figure 2A). Pairwise distances between the 12 conditions were quantified using 1-Spearman’s correlation across all MEG sensors, capturing the unique representational geometry of each electrophysiological signal. Similarly, for the fMRI data, RDMs were generated for each of the 360 HCP cortical parcels, as well as for the single combined empirical MD ROI, by calculating pairwise 1-Spearman’s correlations across all vertices within the respective parcel or ROI (Figure 2B). These RDMs provided a modality-invariant space for evaluating the raw spatial correspondence between electromagnetic and hemodynamic representational patterns.

**Figure 2.**
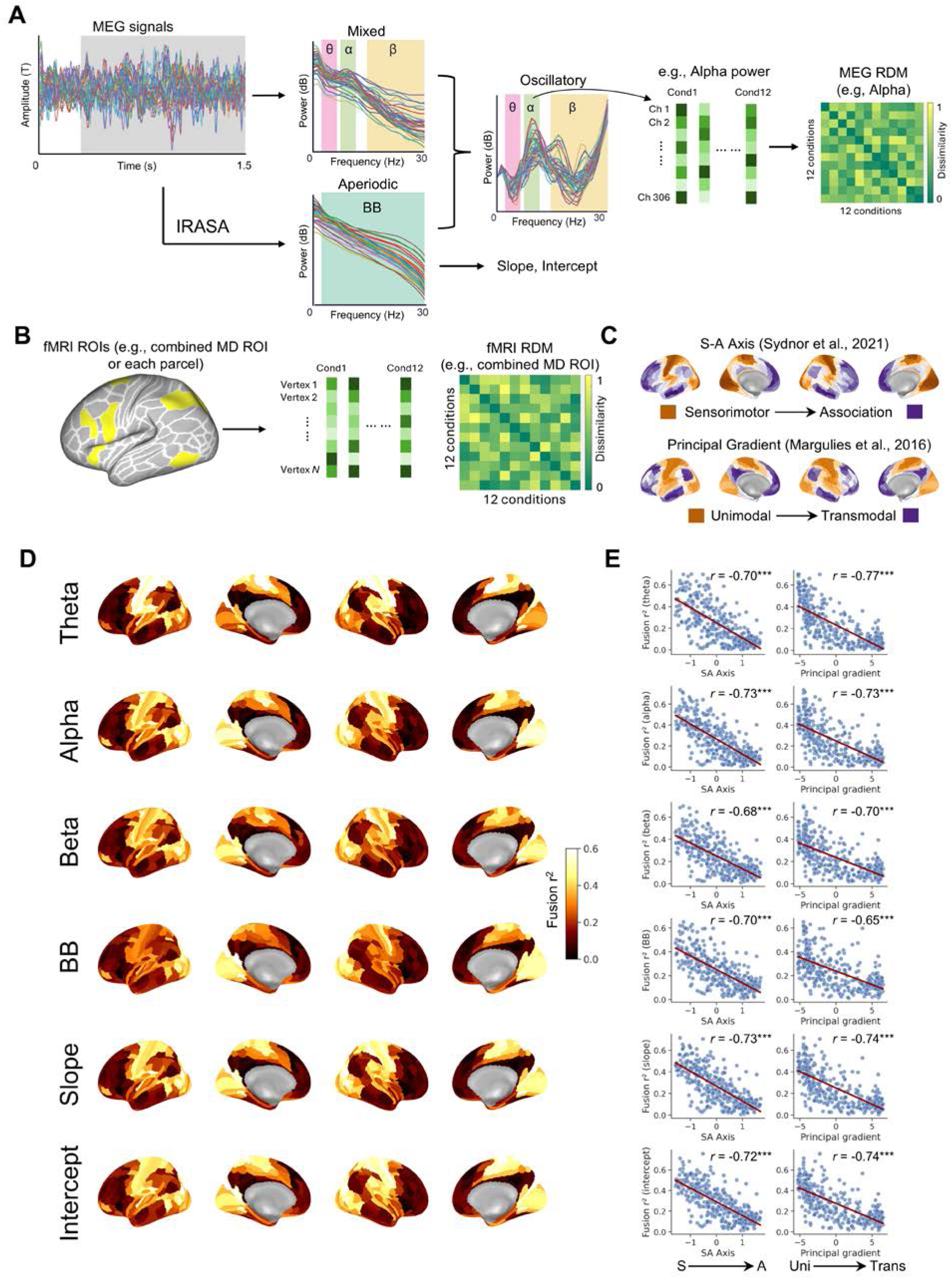
Hierarchical decoupling of raw MEG-fMRI representational correspondence. (A) Construction of MEG RDMs. MEG signals were decomposed using IRASA to separate oscillatory (theta, alpha, beta power) and aperiodic components (BB, slope, and intercept). RDMs for each MEG signal were computed using 1-Spearman’s correlation between 12 conditions (3 tasks × 2 task demands × 2 task contents) across all MEG sensors. We demonstrate this approach with alpha power in the plot. (B) Construction of fMRI RDMs. RDMs were generated for each of the 360 HCP cortical parcels (white), as well as a single combined MD ROI (yellow) encompassing the parcels of the empirical MD network. Vertex-wise fMRI activation patterns were used to compute RDMs for each parcel or ROI using 1-Spearman’s correlation. (C) Canonical macroscale cortical gradients. Spatial distribution of the sensorimotor-association (S-A) axis (top; Sydnor et al. (2021)) and the principal functional gradient (bottom; Margulies et al. (2016)), obtained from the *neuromaps* toolbox (Markello et al., 2022). (D) Whole-brain maps of raw MEG-fMRI fusion strength. For each of the 360 HCP parcels, the raw cross-modal correspondence was quantified as the squared Spearman correlation between the MEG and fMRI RDMs without task-specific theoretical constraints. (E) Spatial relationship with macroscale cortical gradients. Scatter plots illustrate the correlation between the raw fusion strength (from Panel D) and two established cortical hierarchies in Panel C. Each point represents one of the cortical parcels. All *p* values were significant (*** *p*<0.001) after 1000 spatial spin tests.

To ensure the stability of these representational geometries, we evaluated the split-half reliability of the RDMs using a bootstrapping procedure (1000 iterations) coupled with Spearman-Brown correction (see Methods). The MEG representations demonstrated excellent reliability across all electrophysiological signals (all *r*_sb_ > 0.91). Similarly, the fMRI RDMs exhibited high internal consistency, yielding a whole-cortex average reliability of *r*_sb_ = 0.92 across the 360 parcels (see Figure S4 for the whole brain reliability). These highly reliable RDMs provided a robust, modality-invariant space for evaluating the raw spatial correspondence between electromagnetic and hemodynamic representational patterns.

### Hierarchical decoupling of raw MEG-fMRI correspondence alongside the S-A axis

We first conducted a model-free RSA fusion across 360 cortical parcels to characterize the intrinsic spatial relationship between electrophysiological and hemodynamic representations. This analysis quantified the raw correspondence (squared Spearman’s *r*) between the RDMs of each MEG signal and fMRI parcel, without applying any task-specific theoretical constraints.

We observed marked spatial heterogeneity in the strength of MEG-fMRI correspondence across the cortex (Figure 2D). For all six MEG signals (oscillatory and aperiodic), the highest correspondence was consistently localized in unimodal sensorimotor regions, such as the primary visual and somatosensory cortices. In contrast, transmodal association areas, including the frontoparietal MD network and the default network, exhibited weaker raw cross-modal similarity.

To quantify this observation, we performed spatial correlation analyses between the whole-brain fusion maps and two established macroscale gradients: the S-A axis (Sydnor et al., 2021) and the principal functional gradient (Margulies et al., 2016) (Figure 2C). As shown in Figure 2E, we found the raw fusion strength for all MEG signals was significantly and negatively correlated with both the S-A axis (all *r*s < -0.68, *p*s < 0.001) and the principal gradient (all *r*s < -0.65, *p*s < 0.001), after controlling for spatial autocorrelation using spin tests (Vasa & Misic, 2022).

These results demonstrate a clear hierarchical decoupling of raw electro-hemodynamic correspondence during cognitive control tasks: while sensory regions show tight coupling between electromagnetic and hemodynamic representational geometries, these two modalities profoundly diverge in higher-order association cortices including the MD network.

### Demand-related MEG-fMRI fusion in the empirical MD network

To isolate the representational shared variance uniquely driven by specific cognitive components, we employed a model-based commonality analysis framework (Figure 3A). We constructed three theoretical model RDMs to capture within-task demand (easy vs. hard), within-task content (alphanumeric vs. colour), and condition-specific differences in behavioural accuracy (as a control model to regress out variance related to idiosyncratic performance and broad between-task differences). While all three models were included in the analysis to account for overlapping variance, our primary focus was on the demand-specific commonality. In brief, demand-specific commonality quantifies the amount of representational variance shared between the MEG and fMRI signals that can be uniquely explained by task demand, while partialling out the influence of the other models. Crucially, to account for the systematic spatial variation in baseline MEG-fMRI coupling (Figure 2), we utilized relative commonality as our primary metric. This was calculated by normalizing the demand-specific commonality by the total shared variance within each region, thereby ensuring our results reflected the proportion of cross-modal integration dedicated to task demand, independent of regional baseline biases.

**Figure 3.**
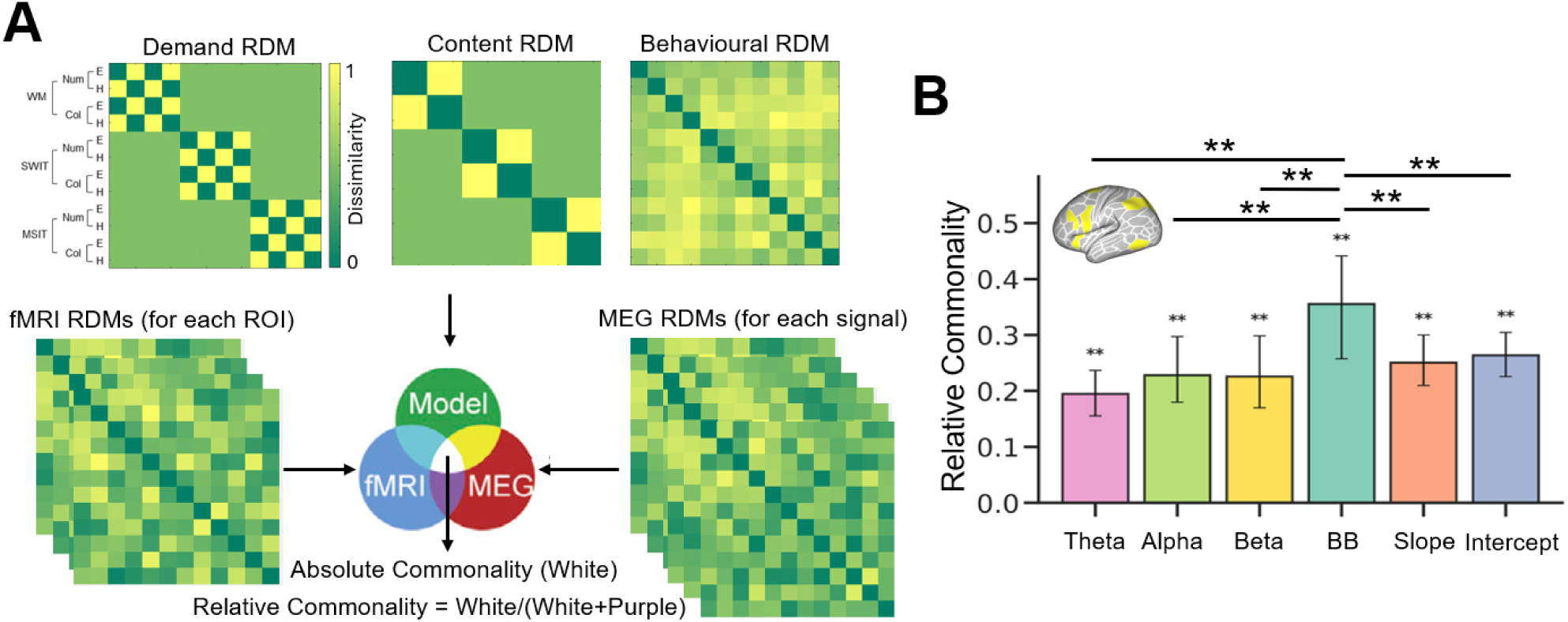
Demand-related MEG-fMRI fusion in the empirical MD network. (A) Model-based fusion approach. To isolate representational shared variance driven by specific cognitive factors, we constructed three model RDMs for capturing within-task demand, within-task content, and behavioural accuracy. The intersection in the Venn diagram represents the unique absolute commonality shared between electromagnetic (MEG) and hemodynamic (fMRI) signals that is explained by a specific model (e.g., task demand) while partialling out the other two models. Relative commonality was calculated by dividing the absolute commonality by the total shared variance between the two modalities (the model-free baseline *R*^2^). (B) Demand-specific relative commonality in the empirical MD network. The bar plot shows the relative commonality (percentage of total cross-modal shared variance) for the six electrophysiological signals within the empirical MD network. Error bars represent 95% confidence intervals derived from 1000 bootstrap iterations. *p*-values were obtained from a permutation-based null distribution, followed by FDR correction (** *p* _FDR_ < 0.01).

Within the empirical combined MD network ROI, we found that task demand captured a significant proportion of the shared variance between electromagnetic and hemodynamic signals, reflecting a robust task-driven representational specialization. While all six MEG signals exhibited significant positive commonality with task demand (all *p*s _FDR_ < 0.01), BB showed the highest relative commonality among them (Figure 3B). Exploratory pairwise comparisons using permutation-based t- tests further revealed that BB’s relative commonality was significantly greater than that of all other oscillatory and aperiodic signals (*p*s _FDR_ < 0.01). These results suggest that BB activity may serve as the primary electrophysiological substrate for the MD network’s response to cognitive demand.

### Demand-specific opposing gradients across the cortical hierarchy

Having identified BB activity as a key substrate for demand representation in the MD network, we next investigated whether this link is locally confined or reflects a broader organizational principle across the association cortex and the whole brain. To do this, we calculated the demand-specific relative commonality in each of the 360 cortical parcels (Figure 4A). Notably, at the whole-brain level, we found the demand-related relative commonality for BB was globally higher than that of all other oscillatory and aperiodic signals. This suggests that broadband aperiodic dynamics provide a globally superior electrophysiological marker for task demand compared to canonical oscillations.

**Figure 4.**
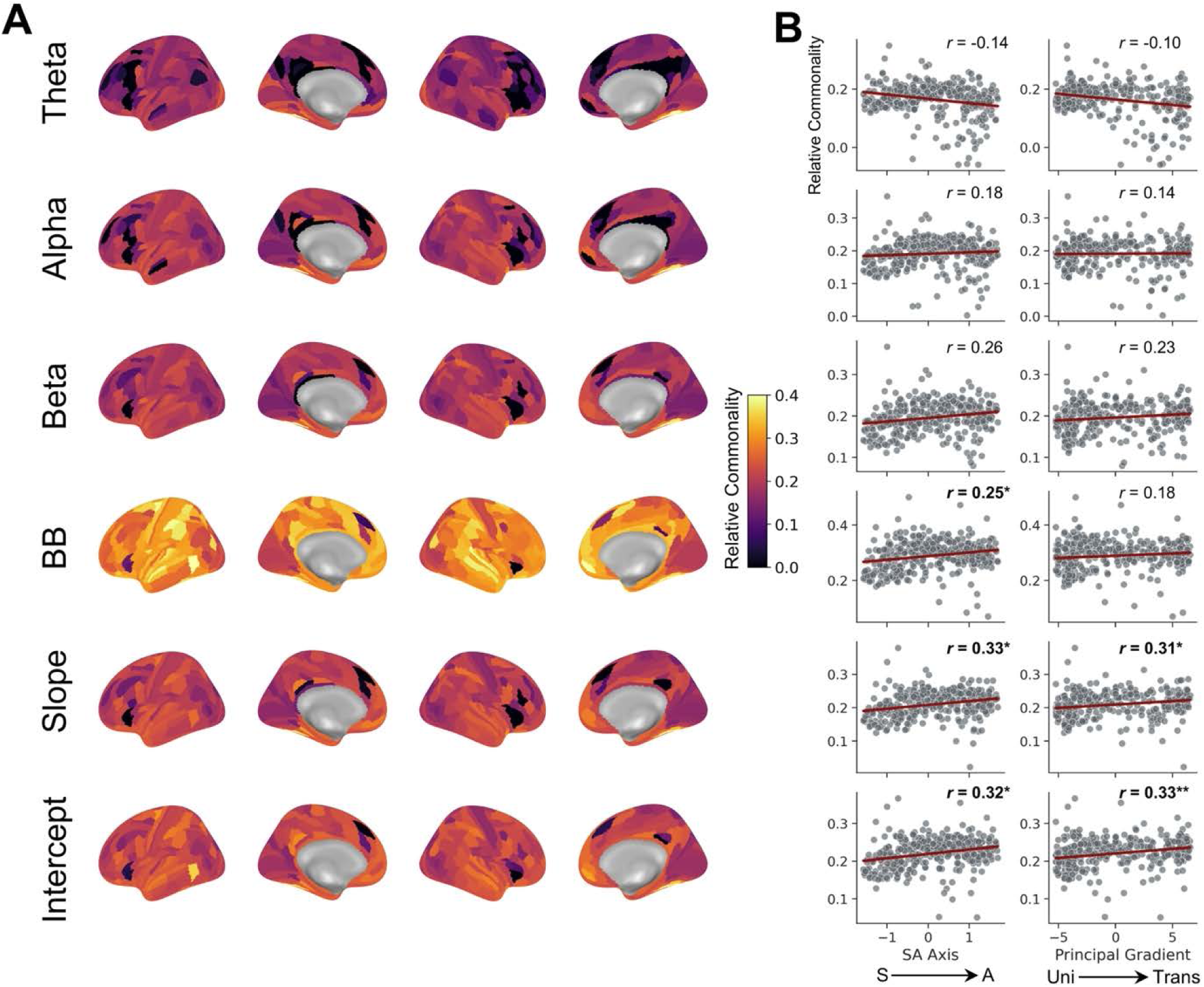
Demand-related MEG-fMRI fusion across the cortical hierarchy. (A) Whole-brain maps of demand-related relative commonality. Demand-specific commonality was projected across the 360 HCP parcels for all six signals. (B) Spatial relationship with macroscale cortical gradients. Scatter plots illustrate the relationship between demand-specific relative commonality and macroscale cortical hierarchies (S-A axis and principal gradient). While raw correspondence was negatively correlated with the S-A axis (see Figure 2), demand-specific commonality for aperiodic components shows a significant positive correlation (* *p* _FDR_ < 0.05, ** *p* _FDR_ < 0.01).

We further examined whether this demand-specific cross-modal commonality follows the macroscale organization of the cortical hierarchy (Figure 4B). In contrast to the hierarchical decoupling observed in our model-free analysis, where raw correspondence was weakest in association areas, the isolation of demand-related information revealed an opposing hierarchical gradient. For aperiodic signals specifically, the demand-specific relative commonality was significantly and positively correlated with the S-A axis (all *r*s > 0.25, *p*s < 0.05) after controlling for spatial autocorrelation using spin tests. Similarly, we found that the aperiodic components (slope and intercept) were significantly and positively correlated with the principal functional gradient (both *r*s > 0.31, *p*s < 0.05). In comparison, no significant correlations were detected between oscillatory components and cortical hierarchy after spin tests.

Collectively, these results reveal that the cross-modal commonality driven by cognitive demand is not locally confined to the MD network, but reflects a broader organizational principle. Notably, although both oscillatory and aperiodic signals showed a comparable negative gradient in raw correspondence, only aperiodic dynamics exhibited a significant demand-related opposing gradient along the S-A axis. This indicates that in higher-order association regions, a greater proportion of the shared electro-hemodynamic variance is dedicated to task-specific information, rather than just reflecting regional variations in baseline coupling strength. These findings position aperiodic activity as the essential bridge that aligns discrete neural modalities into a coherent representational space for cognitive control.

## Discussion

In the present study, we combined fMRI and MEG data across a diverse set of cognitive control tasks to characterize the cross-modal representational architecture of domain-general task demand and its mapping within the domain-general MD network as well as across the cortical hierarchy. We found although raw MEG-fMRI representational correspondence was strongest in unimodal sensorimotor cortex and progressively weakened toward higher-order association cortex, isolating variance uniquely driven by cognitive demand through model-based commonality analysis revealed an opposing hierarchical gradient. Within the MD network, BB emerged as the primary substrate for this cross-modal alignment, providing a more robust bridge for integration than traditional frequency-specific oscillations or other aperiodic indices. Importantly, our whole-brain findings suggest that this demand-driven cross-modal commonality is not strictly confined to the MD network but instead reflects a broader organizational property of the association cortex. Specifically for aperiodic activity, the proportion of cross-modal shared variance attributable to task demand became increasingly prominent along the S-A axis, reaching its peak in the association areas. Collectively, our results position aperiodic dynamics as an essential link that aligns the brain’s hemodynamic and electrophysiological responses into a coherent framework for flexible cognitive control.

Using six subtasks spanning three cognitive control processes, we observation that similar frontal, parietal, and temporal regions are recruited across diverse tasks. This highlights the central role of the MD network in supporting flexible, cross-task cognitive demands. However, compared to the previously defined MD network (Assem et al., 2020), we observed shifts in the precise locations of some MD parcels. For instance, activations around the intraparietal sulcus were more dorsal but less ventral, and the anterior frontal region showed weaker activation during MSIT tasks. These findings may align with prior evidence of task preference within MD regions (Assem et al., 2020; Assem et al., 2024), suggesting that while these broad regions are consistently recruited, their fine-grained activation topography may shift based on task-specific demands. Thus, we think that the MD network may operate as a general and flexible framework, with slightly malleable localization of peak activity reflecting task specificity. Such modulation might reflect the need for MD regions to flexibly interface with different task-specific brain networks, depending on the context.

To examine the intrinsic link between electromagnetic and hemodynamic representational geometries, we first conducted a model-free fusion analysis across the cortex. We found that the raw spatial correspondence between MEG and fMRI signals exhibited marked spatial heterogeneity. Specifically, this raw cross-modal similarity was strongest in unimodal sensorimotor regions and progressively decayed toward transmodal association cortices. This hierarchical decoupling directly aligns with the macroscale organization of the S-A axis (Sydnor et al., 2021) and the principal functional gradient (Dong, Margulies, Zuo, & Holmes, 2021; Huntenburg, Bazin, & Margulies, 2018; Margulies et al., 2016), suggesting that the alignment between fast electrophysiological and slow metabolic signals may be constrained by the brain’s hierarchical scaffold. These results extend the recent observations demonstrating a similar hierarchical decoupling in the resting state (Shafiei et al., 2022), where the functional network architectures of the two modalities systematically converge in unimodal cortex and diverge in transmodal cortex. This spatial heterogeneity is likely rooted in the cortical microarchitecture that the transmodal cortex exhibits distinct laminar differentiation and reduced structure-function coupling compared to primary sensory regions (Huntenburg et al., 2018; Paquola et al., 2019; Suarez, Markello, Betzel, & Misic, 2020; Vazquez-Rodriguez et al., 2019). Crucially, while prior evidence of this transmodal divergence has been largely derived from resting-state paradigms, our findings demonstrate that this intrinsic decoupling is a robust baseline property that persists during active cognitive performance, setting the stage for task-driven mechanisms to bridge this modal gap.

Against this baseline divergence, our model-based fusion analysis reveals a striking representational reconfiguration during active cognitive control. By isolating the variance uniquely attributable to cognitive demand, we demonstrated that the proportion of cross-modal shared variance dedicated to cognitive demand forms an opposing hierarchical gradient. Within the transmodal MD network, BB emerged as the primary electrophysiological signal mediating this cross-modal alignment, significantly outperforming canonical frequency-specific oscillations. This result aligned with our previous MEG findings that BB is the most robust electrophysiological correlate of domain-general cognitive control (Lu et al., 2024). However, rather than being a strictly MD-localized phenomenon, we found this demand-specific cross-modal alignment systematically scales along the S-A axis. This large-scale topography indicates that the integration of electromagnetic and hemodynamic signals may not be a rigid anatomical constraint, but rather a flexible computational resource recruited for high-level cognition.

The superiority of aperiodic activity in bridging this gap provides important mechanistic insights. A growing body of research suggests that aperiodic activity contributes to a wide range of cognitive functions, including working memory, attention, memory consolidation, processing speed, and cognitive control (Donoghue et al., 2020; Gyurkovics, Clements, Low, Fabiani, & Gratton, 2022; He, 2014; Ouyang, Hildebrandt, Schmitz, & Herrmann, 2020; Pei, Northoff, & Ouyang, 2023; Thuwal, Banerjee, & Roy, 2021; Waschke et al., 2021). Unlike rhythmic oscillations, which are typically implicated in cortical gating or the selective routing of information channels (Jensen & Mazaheri, 2010; Vinck, Uran, Dowdall, Rummell, & Canales-Johnson, 2024), aperiodic activity is increasingly recognized as a macroscopic proxy for asynchronous neural firing and global shifts in the E/I balance (Donoghue et al., 2020; Gao et al., 2017; Kałamała et al., 2024; Lu, 2025; Manning, Jacobs, Fried, & Kahana, 2009; Voytek & Knight, 2015). Navigating complex task demands requires transmodal cortices to rapidly modulate these E/I states to optimize network flexibility and integrate diverse computational rules (Waschke et al., 2021). Consequently, our findings suggest that the widespread BOLD activations characterizing cognitive control are deeply tethered to these shifts in asynchronous population-level excitability, positioning aperiodic dynamics as the fundamental neurophysiological currency of cognitive effort.

We acknowledge that the acquisition of fMRI and MEG data from independent participant samples is a limitation of the current study. Due to practical constraints, this design precluded the direct investigation of within-subject cross-modal associations or the influence of individual differences on electro-hemodynamic coupling. However, several lines of evidence validate the robustness of our cross-modal findings. First, our split-half reliability analysis demonstrated that the RDMs for both modalities were highly stable, with reliability coefficients exceeding 0.91 for all MEG signals and reaching 0.92 for the fMRI whole-cortex average. Such internal consistency indicates that our group-level RDMs capture robust, population-level neural signatures rather than noise-driven or sample-specific patterns. Furthermore, the use of a rigorous statistical framework involving both bootstrapping iterations and permutation testing ensures that the observed demand-related representational reconfiguration is a stable phenomenon that generalizes across independent groups. While future studies using simultaneous or within-subject fMRI-MEG recordings would be valuable, the current results provide a reliable characterization of how task demands modulate the alignment of discrete neural modalities across the cortical hierarchy.

In conclusion, our study indicates that aperiodic neural activity serves as an electrophysiological substrate that links electromagnetic and hemodynamic representations of cognitive demand. By applying a multimodal fusion framework, we observed that while overall MEG and fMRI signals exhibit hierarchical decoupling along the S-A axis, the proportion of cross-modal shared variance driven by task demand forms an opposing hierarchical gradient. Within the MD network and higher-order association regions, aperiodic broadband power accounted for a greater proportion of the demand-specific shared cross-modal variance compared to other oscillatory and aperiodic signals, suggesting that it may serve as the primary electrophysiological substrate underlying the MD network’s demand-related activations. These results suggest that the spatial alignment between fast neurophysiological signals and slow metabolic responses is modulated by domain-general task requirements. Collectively, this work identifies aperiodic dynamics as a potential substrate for the large-scale integration of neural representations, providing a framework for understanding how discrete brain imaging modalities converge during active cognitive control.

## Methods

### Participants

Forty-two healthy, right-handed participants were recruited for the fMRI study from the local community and the online participant database (SONA) at the University of Cambridge. Six participants were excluded from the formal fMRI analysis due to behavioural accuracy falling more than three standard deviations below the mean. Consequently, 36 participants (age range: 21–49 years; 20 females, 16 males) were included in the final analysis. For the MEG study, an independent group of 43 participants (age 18-39 years, 31 females and 12 males) underwent formal analysis (see (Lu et al., 2024) for details). All participants for both fMRI and MEG studies were native Mandarin speakers with normal or corrected-to-normal vision and no history of neurological disorders. We obtained written informed consent from all participants, who were compensated for their time. Both studies were approved by the Cambridge Psychology Research Ethics Committee. To control for word-length differences between the alphanumeric and colour WM tasks (i.e., recalling letters and colours) (Baddeley, Thomson, & Buchanan, 1975), we recruited native Mandarin speakers. As explained in (Lu et al., 2024), Mandarin speakers typically encode English letters in English, but process visually presented colours in Mandarin. We reinforced this strategy in our instructions to participants. We selected English letters that are monosyllabic (e.g., “A” /eɪ/, “B” /biː/) and paired them with colours that have monosyllabic names in Mandarin, such as red (/hʊŋ/), yellow (/hwɑːŋ/) and purple (/dzɪ/). This approach ensured that both types of WM stimuli—letters and colours—were encoded as monosyllabic terms, thereby reducing phonetic word length effects and helping to balance cognitive demand between the alphanumeric and colour conditions (Chan & Elliott, 2020; Stigler, Lee, & Stevenson, 1986).

### Task paradigms

To investigate domain-general neural activity associated with cognitive demand across multiple tasks, we employed three cognitive paradigms (a working memory task, WM; a switching task, SWIT; and a multi-source interference task; MSIT). We manipulated each task to have two levels of cognitive demand (hard vs. easy) and two types of task content (alphanumeric vs. colour) (Figure 1). This resulted in six subtasks (3 tasks × 2 content) and 12 unique conditions (3 tasks × 2 demand × 2 content). The fMRI and MEG study designs differed in two aspects. First, in the fMRI study, both task demand and content were blocked to enhance signal-to-noise ratio, whereas in the MEG study, only task content was blocked. Second, in the fMRI study, fixation durations within each trial were reduced to increase the number of trials per 16-second block, thereby optimizing design efficiency. To ensure these design differences did not significantly impact between-modality comparisons, we conducted a pilot study with two participants who each completed two MEG sessions, one using the MEG design and the other following the fMRI design. Results revealed strong positive correlations between flattened condition-by-condition RDMs across all oscillatory and aperiodic responses for both participants (all *p*s < 0.003), confirming that the designs yielded comparable outcomes.

In the fMRI study, each participant completed six runs. Each run consisted of 24 task blocks (12 conditions × 2 repetitions) and 12 fixation blocks, with each block lasting 16 seconds. Easy and hard blocks of one subtask were paired (easy followed by hard, or hard followed by easy) and the subtask order was counterbalanced across runs and subjects. A fixation block (16 s) followed every 2 paired task blocks. At the start of each paired subtask, a 4-second cue was displayed on the screen to prepare participants. Task procedures for the MEG study are detailed in (Lu et al., 2024).

For the WM task, we employed a modified Sternberg task (Sternberg, 1966). In the fMRI study, each trial began with a fixation cross displayed for 0.5 ± 0.1 s (1.5 ± 0.1 s in the MEG study), followed by the presentation of four items (letters for the alphanumeric subtask and coloured circles for the colour subtask) arranged horizontally at the centre of the screen for 0.3 seconds. In the hard condition, the memory set consisted of four unique items, while the set in the easy condition consisted of two unique items flanked by a # (alphanumeric subtask) or black circles (colour subtask). Thus, the physical size and the visual content were similar in the two conditions. After a 1.5 s delay (2 s in the MEG study), a probe appeared at the centre of the screen and the participant were required to press a button within 1.5 s (2 s for MEG) to indicate whether the probe was part of the memory set or not (left button for ’yes,’ right button for ’no’). We used letters from the list “B”, “D”, “G”, “K”, “P”, “Q”, and “R”, and selected the colours from red, orange, yellow, green, blue, purple, and pink.

For the SWIT task, after a fixation cross was shown for 0.5 ± 0.1 s (1.5 ± 0.1 s for MEG), one item (a two-digit number for the alphanumeric subtask and a coloured circle or triangle in the colour subtask) was presented at the centre of the screen within a square or a diamond for 3 s. For the alphanumeric subtask, if the stimulus was surrounded by a square, participants were instructed to indicate whether the number was odd or even (left button for odd, right button for even). If the stimulus was surrounded by a diamond, participants were required to indicate whether the number was divisible by 3 (left button for yes, right button for no). For the colour subtask, participants were required to indicate whether the stimulus was blue or red if the stimulus was surrounded by a square (left button for red, right button for blue), and to indicate whether the stimulus was a circle or a triangle if the stimulus was surrounded by a diamond (left button for circle, right button for triangle). Switch trials (the rule for the present trial was different from the last trial) were considered as the hard condition and repeat trials (the rule for the present trial repeated the last trial) were considered as the easy condition. The stimuli were selected from 12, 13, 15, and 16 for the alphanumeric subtask and selected from blue circle, red circle, blue triangle, and red triangle for the colour subtask.

The MSIT task served as an inhibitory control task. Each trial began with a fixation cross displayed for 0.5 ± 0.1 s (1.5 ± 0.1 s in the MEG study), followed by a row of three items (digits for the alphanumeric subtask and coloured circles for the colour subtask) presented at the centre of the screen for 3 seconds. Participants were required to identify the unique target item among the three and to press a button with one of three fingers within 3 s. The target item was “1” (left response), “2” (middle), or “3” (right) for the alphanumeric subtask and was “red” (left), “green” (middle), or “blue” (right) for the colour subtask. Participants underwent a practice to match “1”, “2”, and “3” (or “red”, “green”, and “blue”) to three buttons. In the hard (incongruent) condition, the target item was presented in the position incongruent with the required response (e.g., “1” or “red” presented at the third position), and always flanked by different interfering numbers or colours (e.g., “331” or “blue, blue, red”). In the easy (congruent) condition, the target item was presented in the position compatible with the required response and flanked by 0 or “X” in the alphanumeric subtask (e.g., “100” or “1XX”) while flanked by black circles in the colour subtask (e.g., “red, black, black”).

### MRI data acquisition

MRI data were collected using a 3 T Siemens Prisma scanner with a 32-channel RF head coil. We use the MRI Connectome Coordination Facility (CCF) acquisition protocols for the HCP Young Adult cohort (https://protocols.humanconnectome.org/CCF/; package date 14 July 2016). These protocols are largely consistent with those described in previous HCP studies (Glasser et al., 2013; Ugurbil et al., 2013), with some minor variations. Each participant underwent both structural and functional MRI scans within a single session. Structural imaging included one T1w MPRAGE and one T2w SPACE scan, each acquired at 0.8 mm isotropic resolution. Task fMRI data were collected across six runs (∼60 min in total) using a whole-brain, multi-band gradient echo planar imaging (EPI) sequence with 2.4 mm isotropic resolution (TR = 1.13 s, TE = 37 ms, multi-band factor = 4). Task EPI runs were acquired in pairs of reversed phase-encoding directions (AP/PA). Spin echo phase reversed images in the anteroposterior directions (AP/PA) matched to the gradient echo fMRI images were acquired at the beginning and middle of functional scanning sessions to (i) correct T1w and T2w images for readout distortion to enable accurate T1w to T2w registration, (ii) enable accurate cross-modal registrations of the fMRI images to the T1w image in each subject, (iii) compute a more accurate fMRI bias field correction and (iv) segment regions of gradient echo signal loss.

### MRI data preprocessing

MRI data preprocessing closely followed the HCP’s minimal preprocessing pipelines (Glasser et al., 2013) as detailed in previous studies(Assem, Shashidhara, Glasser, & Duncan, 2022; Assem et al., 2024). The preprocessing utilized HCP pipelines versions 4.0.0 (scripts available at: https://github.com/Washington-University/HCPpipelines). Below is a brief overview, with specific steps and differences noted.

For each subject, structural images (T1w and T2w) were used to extract cortical surfaces and segmentation of subcortical structures. Functional images were mapped from volume to surface space and combined with subcortical data in volume to form the standard CIFTI grayordinates space. We smoothed the data by a 2mm FWHM kernel in the grayordinate space, which avoids mixing data across gyral banks for cortical surface data and avoids mixing data across major structure borders for subcortical data.

Task fMRI data were further cleaned for spatially specific noise using spatial independent component analysis-based Xnoiseifier (ICA+FIX (Salimi-Khorshidi et al., 2014)). ICA+FIX was applied to concatenated task runs. An improved FIX classifier was used (HCP_Style_Single_Multirun_Dedrift in ICAFIX training folder) for more accurate classification of noise components in task fMRI datasets. For accurate cross-subject registration of cortical surfaces, the multimodal surface matching (MSM) algorithm was applied. Initially, “sulc” cortical folding maps were gently registered in the MSMSulc registration, optimizing for functional alignment without overfitting folds. Subsequently, a combination of myelin maps and the task fMRI data were used to functionally align the data (MSMAll; (Robinson et al., 2018; Robinson et al., 2014)).

### Task fMRI analysis

We performed task fMRI analysis using HCP pipelines version 4.0.0, following the default steps described in (Barch et al., 2013). Briefly, autocorrelation was estimated on the cortical surface using FSL’s FILM with default parameters from the HCP task fMRI analysis scripts. Activation estimates were computed for the preprocessed functional time series of each run using a general linear model (GLM) implemented in FSL’s FILM (Woolrich, Ripley, Brady, & Smith, 2001).

For both univariate activation analysis and RSA-based fusion analysis, we included 12 regressors in the GLM, corresponding to 3 tasks × 2 content types (Colour vs. Alphanumeric) × 2 demand levels (hard vs. easy). Each predictor had a unitary height and covered the entire block duration (16 s, from block onset to the offset of the final trial). Fixation blocks were not explicitly modelled and therefore contributed to the implicit baseline. All regressors were then convolved with a canonical hemodynamic response function and its temporal derivative. The time series and the GLM design were temporally filtered with a Gaussian-weighted linear highpass filter with a cutoff of 200 s. Finally, the time series was prewhitened within FILM to correct for autocorrelations. Surface-based autocorrelation estimate smoothing was incorporated into FSL’s FILM at a sigma of 5 mm.

Percent signal change (i.e., normalised beta estimates) was calculated as follows: 100×(beta/10,000), where 10,000 corresponds to the mean scaling of each vertex time series during pre-processing. For RSA-based fusion analysis, we used condition-specific beta estimates (i.e., condition-specific activation relative to the implicit baseline). These percentage signal change values were then used for subsequent fusion analyses. For univariate activation analysis, fixed-effects analyses were conducted using FSL’s FEAT to estimate the average effects across runs within each subject. We defined contrasts to assess task demand (hard > easy) for each subtask. For cortical parcellation, the group-average HCP multimodal parcellation (MMP1.0) was used (Glasser et al., 2016).

### Definition of the empirical MD ROI

To maximize the spatial sensitivity of our fusion analyses to the specific cognitive demands of the current paradigms, we defined an empirical MD ROI based on the regions showing the strongest activation in the hard > easy contrast. Given that activation patterns were highly consistent across hemispheres, we first averaged the beta estimates across corresponding regions in the left and right hemispheres to improve the signal-to-noise ratio. For each participant, we then averaged the vertex-wise beta values within each of the 180 cortical areas defined by the HCP-MMP1.0 parcellation (Glasser et al., 2016).

To identify regions consistently recruited by task demand across different cognitive domains, we performed a group-level analysis for each of the six subtask contrasts (3 tasks × 2 stimulus contents). For each contrast, we identified areas where the hard > easy activation was significantly greater than zero across participants (*p* < 0.05, Bonferroni corrected for 180 areas). Areas that reached this significance threshold in at least five out of the six subtask contrasts were selected for the empirical MD ROI.

This procedure resulted in a set of 22 cortical parcels: 6a, PEF, IFJp, 6r, IFSp, p9-46v, AVI, FOP4, a32pr, SCEF, AIP, LIPd, IPS1, IP0, IP1, IP2, 7PL, 7Am, 7Pm, MIP, VIP, and PH. These empirical MD patches closely aligned with the predefined MD patches described in previous literature (Assem et al., 2020), with approximately half of the areas overlapping with the predefined extended MD network and the remaining half located in immediately adjacent cortical regions.

### MEG data acquisition and processing

As described in (Lu et al., 2024), MEG data were collected in a magnetically shielded room with a Neuromag Vectorview system (Elekta AB, Stockholm, Sweden) with 306 channels (204 planar gradiometers and 102 magnetometers). While our original MEG study (Lu et al., 2024) simultaneously recorded MEG and EEG, only MEG channels were analysed here. We focused exclusively on sensor-level MEG data without source-level reconstruction.

Following preprocessing steps described in (Lu et al., 2024), we downsampled the MEG data to 200 Hz. After removing event-related fields (i.e., the average signal across trials) from each trial for each condition (Cohen & Donner, 2013), we used irregular resampling auto-spectral analysis (IRASA) (Wen & Liu, 2016b) to separate the oscillatory and the aperiodic components from the mixed power spectrum. We performed IRASA over the frequency range of 1–35 Hz (1 Hz steps) using time windows from 0.3 to 1.5 seconds relative to stimulus onset for each subtask (detailed IRASA settings are provided in (Lu et al., 2024)). This procedure yielded distinct aperiodic and oscillatory components, which were used to calculate oscillatory and aperiodic responses of interest (oscillatory: theta, alpha, and beta power; aperiodic: broadband power, slope, and intercept). For oscillatory signals of interest, we averaged the power for each frequency band using the ranges theta: 3-7 Hz; alpha: 8-12 Hz; and beta: 15-30 Hz. For aperiodic broadband power (BB), we averaged the aperiodic spectrum power across 3-30 Hz. To obtain the slope and intercept of the aperiodic components, we used least squares estimation to fit a linear function to the estimated aperiodic power spectrum in log-log coordinates, extracting the slope and the intercept for later analyses.

### MEG-fMRI fusion analysis

We employed a two-level MEG-fMRI fusion framework, incorporating both model-free and model-based RSA analysis. This approach utilizes RDMs to abstract information across imaging modalities, enabling direct comparisons of neural representational patterns between MEG and fMRI data (Cichy & Oliva, 2020; Kriegeskorte, Mur, & Bandettini, 2008).

### RDM construction

The RDMs consisted of 12 × 12 matrices, representing all combinations of task (3) × task demand (2) × task content (2). Each cell in the RDM (dissimilarity) was quantified as 1-Spearman’s *r* between condition pairs. For the sensor-space MEG data, we constructed an RDM for each signal of interest (oscillatory: theta, alpha, and beta power; aperiodic: BB, slope, and intercept). To obtain robust estimates of the representational geometries, we employed a bootstrapping procedure with 1000 iterations, where MEG participants were randomly sampled with replacement to calculate group-averaged MEG RDMs in each iteration.

For the fMRI data, we constructed RDMs for the empirical combined MD ROI and the 360 HCP cortical parcels. fMRI dissimilarities were calculated based on normalized beta estimates (100 × [beta/10,000]) across all vertices within each parcel/ROI. Unlike the MEG data, fMRI RDMs were derived from the fixed group-averaged data across all participants to serve as a stable spatial reference for the fusion.

We constructed three model RDMs to isolate specific representational components: task demand, task content, and behavioural accuracy. For the demand and content models, condition pairs with the same feature within a task were coded as 0 (similar) and different features as 1 (dissimilar). Cross-task similarities were set to 0.5 to account for large inherent differences between tasks. To control for irrelevant between-task variability, we also included a behavioural RDM calculated by 1-Spearman’s *r* of behavioural accuracy scores across participants. This behavioural model was averaged across MEG and fMRI sessions due to their high correlation (*r* = 0.73, *p* < 0.001). Notably, the behavioural RDM showed negligible correlation with the full demand model (*r* = 0.04, *p* = 0.77), indicating that our demand-specific commonality reflects conceptual task requirements rather than idiosyncratic performance variability.

#### Split-half reliability analysis

To evaluate the stability and generalizability of the neural representational geometries, and to ensure that the representational patterns were not driven by subject-specific noise within our independent samples, we conducted a split-half reliability analysis for both the MEG and fMRI datasets.

For the MEG data, we implemented a resampling procedure with 1000 iterations. In each iteration, the sample of MEG participants was randomly divided into two equal and independent halves. Group-average RDMs were then constructed for each half across the six electrophysiological signals (theta, alpha, beta, BB, slope, and intercept). We quantified the internal consistency of the representational geometry by calculating Spearman’s correlation between the upper-triangular elements of the two group-average RDMs.

An identical procedure was applied to the fMRI data. Across 1000 random iterations, the fMRI participants were split into two independent halves, and group-average RDMs were computed for each of the 360 cortical parcels defined by the HCP-MMP1.0 parcellation. The regional reliability was similarly assessed using the Spearman correlation between the two split-half RDMs for each parcel.

Because splitting the data reduces the sample size by half, the resulting correlation systematically underestimates the true reliability of the full sample. To correct for this attenuation, we applied the Spearman-Brown prediction formula to the correlation coefficient obtained in each iteration: *r*_sb_ = 2*r*/(1+*r*). The final reliability metric for each MEG signal and each fMRI cortical parcel was defined as the mean of the Spearman-Brown corrected correlation coefficients *r*_sb_ across all 1000 iterations. This provided a robust estimate of the upper bound of the representational signal-to-noise ratio within each modality.

#### Model-free fusion

To investigate the baseline electro-hemodynamic coupling, we conducted a model-free fusion by extracting the upper triangular portions of the group-averaged MEG RDMs (across six MEG signals of interest) and fMRI RDMs (across 360 parcels) and calculating the squared Spearman’s rank correlation coefficient between them.

#### Model-based fusion

To isolate the shared neural representations uniquely driven by task demand, we performed a model-based commonality analysis (Hebart & Baker, 2018). As described in the RDM construction, this analysis was embedded within a bootstrapping procedure (1000 iterations) to obtain robust group-level estimates and confidence intervals. In each bootstrap iteration, for each oscillatory or aperiodic signal and for each parcel/ROI, we used commonality analysis to estimate the shared variance between three RDMs: the group-averaged MEG RDM, the fMRI RDM, and the model RDM. For each MEG signal, parcel/ROI, and model, we calculated the difference between two squared semipartial correlation coefficients using Spearman correlation. Both semipartial correlation coefficients reflected the proportion of variance shared between MEG and fMRI, with the first coefficient partialling out all three models from the MEG RDM, the second partialling out all models except the model of interest from the MEG RDM. By comparing the variance when our model of interest was included or not, we obtained a measure of the variance shared between MEG and fMRI that can be uniquely explained by each model. In this way we computed the model fit for each MEG signal per parcel/ROI. The commonality *C* between MEG signal *f* and parcel/ROI *j* can be formally described as:

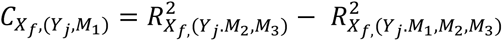

where *X* reflects MEG signals, *Y* reflects fMRI signals, *M_1_* reflects the model of interest (e.g., the demand model), and *M_2_* and *M_3_* reflect the models that are partialled out (e.g., the content and behavioural accuracy models).

To account for the substantial regional variations in baseline MEG-fMRI coupling strength across the cortical hierarchy, we defined relative commonality as our primary metric. Relative commonality was calculated by dividing the absolute commonality *C* by the total shared variance between the two modalities (i.e., the model-free baseline *R*^2^ for that specific MEG component and parcel/ROI).

### Cortical gradients and spatial analysis

To investigate the relationship between cross-modal coupling and macroscale brain organization, we conducted spatial correlation analyses between our fusion maps and canonical cortical hierarchies. We used the *neuromaps* Python toolbox (Markello et al., 2022) to obtain two representative macroscale hierarchies: the sensorimotor-association (S-A) axis (Sydnor et al., 2021) and the principal gradient of functional connectivity (Margulies et al., 2016). Prior to calculating the spatial correlations, we applied a robust outlier detection procedure based on the median absolute deviation (MAD) to ensure that the results were not driven by extreme values or localized artifacts in specific parcels. For each cortical map, we calculated the MAD, defined as the median of the absolute deviations from the data’s median (MAD = median(|*x_i_*-median(X)|)). Parcels with values exceeding 5 times the MAD from the median were identified as extreme outliers and excluded from the subsequent spatial correlation analysis. Finally, we calculated the spatial Spearman rank correlation coefficients between the remaining parcels of each fusion map and the target macroscale hierarchies.

### Statistics

To generate the thresholded activation map for the hard > easy contrast across tasks, we first conducted one-sample t-tests against 0 (right-tailed) for each vertex of the averaged activation map, resulting in a whole-brain t-value map. We then applied Bonferroni correction across all cortical vertices (corrected threshold: *p* = 8.42×10^-7^) and only displayed vertices with *p* values below this threshold.

For the fusion analysis, we computed the group-level RDMs using a bootstrapping procedure for the MEG signals and a fixed group average for the fMRI parcels/ROI. This resulted in a group-level distribution of commonality values for each MEG signal and each fMRI parcel/ROI. Because these metrics were derived at the group level, traditional random-effects statistics across participants were not applicable. Instead, we employed permutation testing to assess statistical significance. To estimate the null distribution, we performed 1000 permutations by randomly shuffling the rows and column labels of the group-average fMRI RDM while maintaining its internal structure (i.e., simultaneously permuting rows and columns). This approach disrupts the alignment between fMRI and MEG representations while preserving the inherent covariance structure of the data.

For each permutation, we recalculated the model-based MEG-fMRI fusion for each MEG signal using the permuted fMRI matrices. The significance of the observed commonality was determined by comparing the mean of the bootstrapped observed values to the null distribution from the permutations, yielding one-tailed *p*-values. False discovery rate (FDR) correction was applied across the six MEG signals for each parcel/ROI and each model. To test for commonality differences between MEG signal pairs (e.g., BB vs. alpha) per parcel/ROI, we compared the observed difference in mean commonality values to a null distribution of differences generated from the permutations. For each MEG signal, FDR correction was applied across the five other MEG signals for each parcel/ROI and model.

When evaluating the spatial correlation between whole-brain cortical maps, traditional degrees of freedom assumptions are invalid due to the high spatial autocorrelation among neighbouring cortical regions. Therefore, we employed a strict spatial permutation framework (spin tests; Vasa and Misic (2022) with 1000 rotations to determine statistical significance. Specifically, we mapped the centroid coordinates of the 360 HCP parcels to the *fsaverage* spherical surface and applied 1000 random 3D rotations. A KD-Tree nearest-neighbor algorithm was used to reassign the rotated coordinates to the corresponding cortical parcels, effectively decoupling the spatial alignment between the target macroscale gradients and our fusion maps while preserving their inherent topological autocorrelation. We computed the *p*_spin_ values by comparing the empirically observed spatial Spearman’s correlations against the null distribution generated from these 1,000 spatial spins. Finally, FDR correction was applied across the *p*_spin_ values of the six MEG signals within each target gradient map.

## Acknowledgements

This project was supported by UKRI MRC intramural funding (MC_UU_00030/15) to A.W., a Gates Cambridge Scholarship (OPP1144) and a postdoctoral fellowship from the Canadian Institutes for Health Research (200883) awarded to R.L. For the purpose of open access, the author has applied a Creative Commons Attribution (CC BY) licence to any Author Accepted Manuscript version arising from this submission.

## Author Contributions

R.L. and A.W. conceived the project; R.L., M.A., J.D., and A.W. designed the experiment; R.L., M.A., and A.W. implemented the experiment; R.L. and M.A., conducted the experiment; R.L. and M.A analysed data; R.L. wrote the first draft of the paper; All authors contributed to the final draft of the paper; A.W. provided overall supervision; and A.W. and R.L contributed funding.

## Supplementary

**Table S1.** Descriptive statistics (fMRI study): Mean and standard deviation (in brackets) of response error and response time (RT), N = 36.

|  |  | WM |  | SWIT |  | MSIT |  |
| --- | --- | --- | --- | --- | --- | --- | --- |
|  |  | Easy | Hard | Easy | Hard | Easy | Hard |
| Accuracy (%) | Alphanumeric | 95.56 (5.02) | 88.26 (7.41) | 96.70 (3.20) | 95.06 (3.98) | 99.72 (0.45) | 97.67 (2.52) |
|  | Colour | 94.98 (4.84) | 83.93 (9.20) | 96.47 (4.22) | 94.59 (4.76) | 99.49 (0.76) | 96.35 (3.73) |
| RT (ms) | Alphanumeric | 785 (111) | 903 (115) | 965 (161) | 1278 (181) | 607 (109) | 874 (124) |
|  | Colour | 728 (104) | 847 (126) | 915 (170) | 1271 (190) | 620 (121) | 932 (142) |

**Figure S1.**
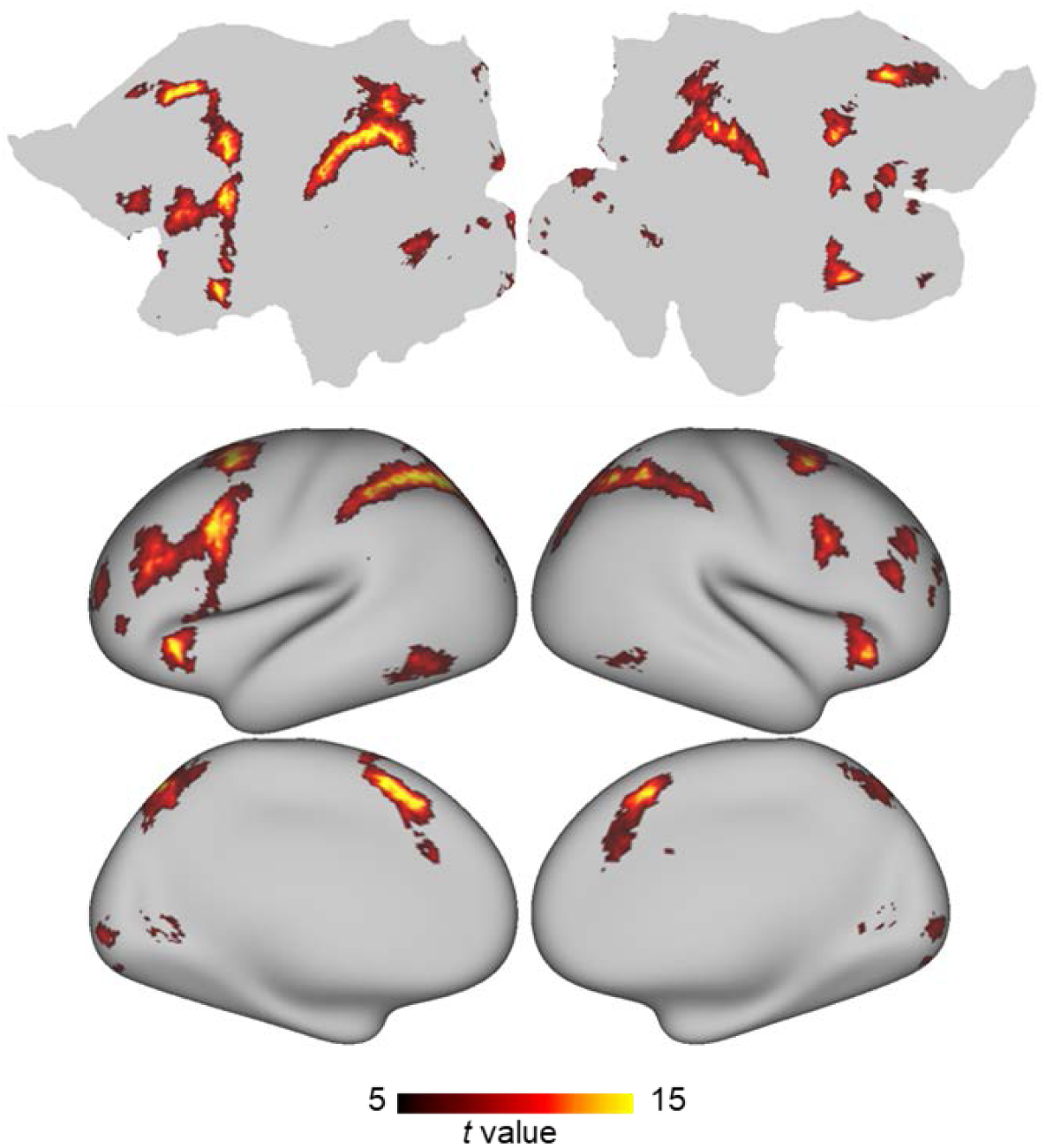
Thresholded t-value maps for hard > easy contrast across subtasks. Thresholded t-value maps computed from activation maps averaged across six hard>easy contrasts [3 tasks (WM, SWIT, and MSIT) × 2 contents (alphanumeric and colour)]. Results were thresholded using Bonferroni correction across all cortical vertices (corrected threshold: *p* = 8.42 × 10⁻⁷), with only significant vertices shown.

**Figure S2.**
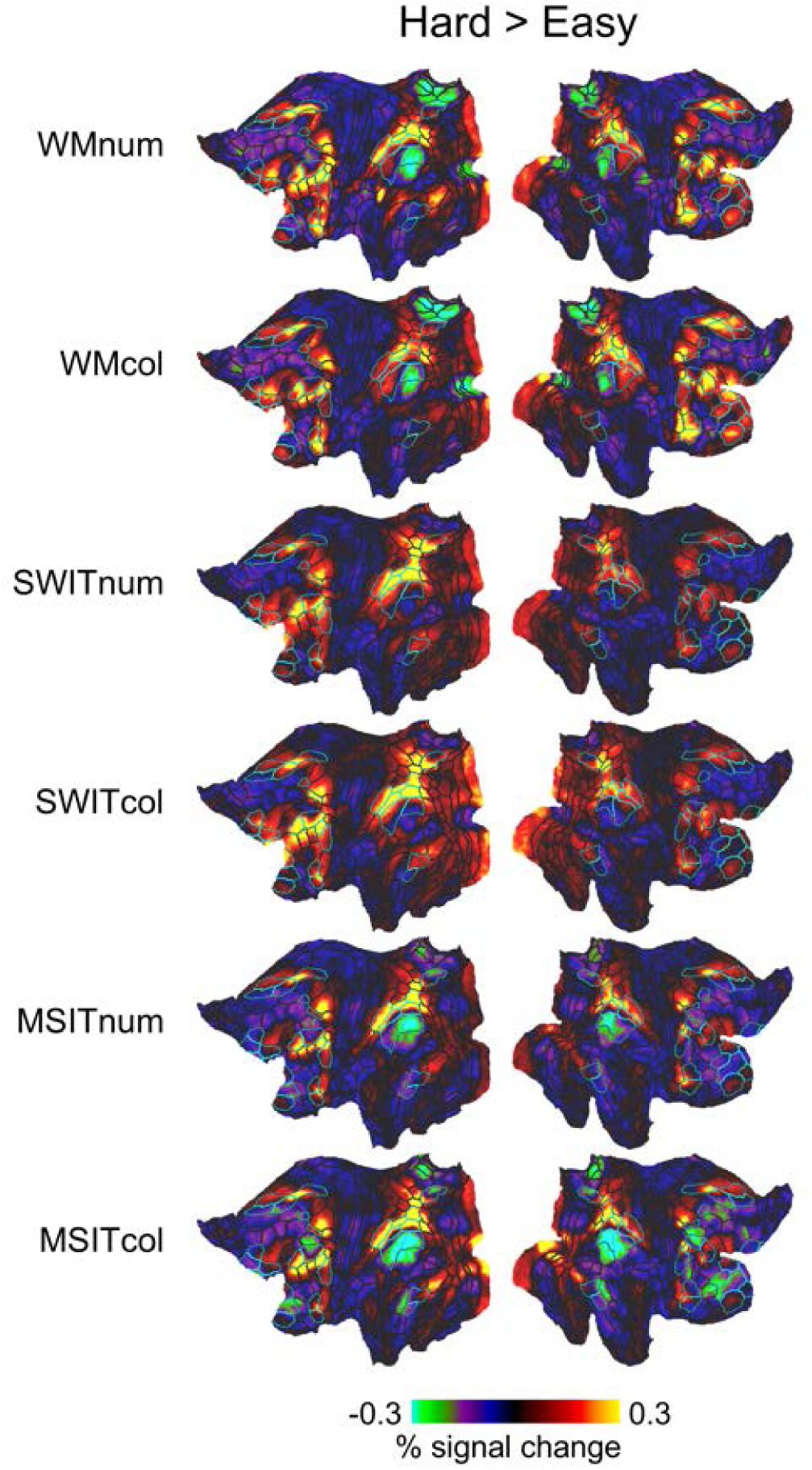
Univariate activation maps for each subtask. Group average activation maps for the hard > easy contrast for each subtask. WM: Working memory task; SWIT: Switching task; MSIT: Multi-source interference task; num: Alphanumeric task; col: Colour task. Cyan borders surround the extended MD areas defined in (Assem et al., 2020).

**Figure S3.**
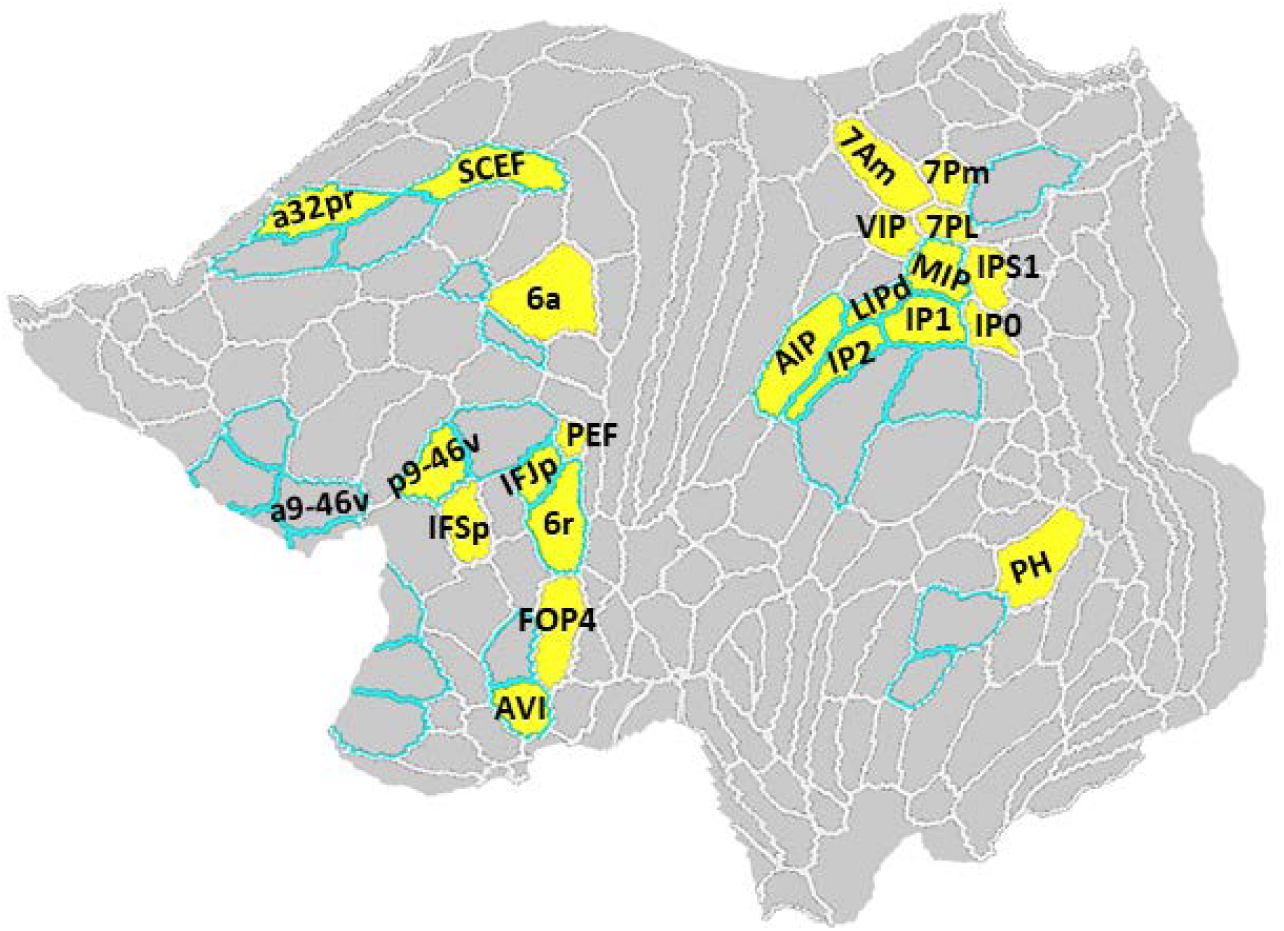
Empirical MD patches defined in this study. Empirical MD patches (yellow) identified using the current dataset. The cortical surface was parcellated into 360 regions (180 per hemisphere) based on the HCP-MMP1.0 Parcellation (outlined in white) (Glasser et al., 2016). Cyan borders surround the extended MD areas defined in (Assem et al., 2020).

**Figure S4.**
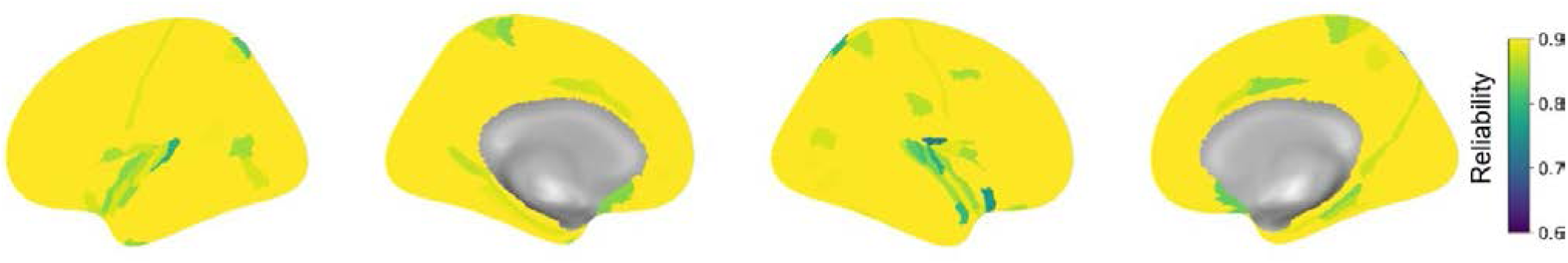
Whole-brain split-half reliability of fMRI representational geometries. The cortical maps illustrate the spatial distribution of the split-half reliability for the fMRI RDMs across the 360 HCP parcels. Reliability was estimated using a bootstrapping procedure with 1000 iterations, where the fMRI participant sample was randomly divided into two independent halves in each iteration. The regional reliability for each parcel was quantified as the Spearman rank correlation between the group-average RDMs of the two halves, followed by the Spearman-Brown correction.

## References

Assem, M., Glasser, M. F., Van Essen, D. C., & Duncan, J. (2020). A Domain-General Cognitive Core Defined in Multimodally Parcellated Human Cortex. Cereb Cortex, 30(8), 4361–4380. doi:10.1093/cercor/bhaa023

Assem, M., Shashidhara, S., Glasser, M. F., & Duncan, J. (2022). Precise Topology of Adjacent Domain-General and Sensory-Biased Regions in the Human Brain. Cereb Cortex, 32(12), 2521–2537. doi:10.1093/cercor/bhab362

Assem, M., Shashidhara, S., Glasser, M. F., & Duncan, J. (2024). Basis of executive functions in fine-grained architecture of cortical and subcortical human brain networks. Cereb Cortex, 34(2). doi:10.1093/cercor/bhad537

Baddeley, A., Thomson, N., & Buchanan, M. (1975). Word length and the structure of short-term memory. Journal of Verbal Learning and Verbal Behavior, 4(6), 575–589.

Barch, D. M., Burgess, G. C., Harms, M. P., Petersen, S. E., Schlaggar, B. L., Corbetta, M.,…Consortium, W. U.-M. H. (2013). Function in the human connectome: task-fMRI and individual differences in behavior. Neuroimage, 80, 169–189. doi:10.1016/j.neuroimage.2013.05.033

Baum, G. L., Cui, Z., Roalf, D. R., Ciric, R., Betzel, R. F., Larsen, B.,…Satterthwaite, T. D. (2020). Development of structure-function coupling in human brain networks during youth. Proc Natl Acad Sci U S A, 117(1), 771–778. doi:10.1073/pnas.1912034117

Bender, A., Zhao, C., Vogel, E., Awh, E., & Voytek, B. (2025). Differential representations of spatial location by aperiodic and alpha oscillatory activity in working memory. Proc Natl Acad Sci U S A, 122(30), e2506418122. doi:10.1073/pnas.2506418122

Bernhardt, B. C., Smallwood, J., Keilholz, S., & Margulies, D. S. (2022). Gradients in brain organization. Neuroimage, 251, 118987. doi:10.1016/j.neuroimage.2022.118987

Chan, M. E., & Elliott, J. M. (2020). Cross-Linguistic Differences in Digit Memory Span. Australian Psychologist, 46(1), 25–30. doi:10.1111/j.1742-9544.2010.00007.x

Chikhi, S., Matton, N., & Blanchet, S. (2022). EEG power spectral measures of cognitive workload: A meta-analysis. Psychophysiology, 59(6), e14009. doi:10.1111/psyp.14009

Cichy, R. M., & Oliva, A. (2020). A M/EEG-fMRI Fusion Primer: Resolving Human Brain Responses in Space and Time. Neuron, 107(5), 772–781. doi:10.1016/j.neuron.2020.07.001

Cohen Kadosh, R. (2025). Rethinking excitation/inhibition balance in the human brain. Nat Rev Neurosci, 26(8), 451–452. doi:10.1038/s41583-025-00943-0

Cohen, M. X., & Donner, T. H. (2013). Midfrontal conflict-related theta-band power reflects neural oscillations that predict behavior. J Neurophysiol, 110(12), 2752–2763. doi:10.1152/jn.00479.2013

Cole, M. W., & Schneider, W. (2007). The cognitive control network: Integrated cortical regions with dissociable functions. Neuroimage, 37(1), 343–360. doi:10.1016/j.neuroimage.2007.03.071

Dixon, M. L., Vega, A. D. L., Mills, C., Andrews-Hanna, J., Spreng, R. N., Cole, M. W., & Christoff, K. (2018). Heterogeneity within the frontoparietal control network and its relationship to the default and dorsal attention networks. Proc Natl Acad Sci U S A, 115(13), E3068. doi:10.1073/pnas.1803276115

Dong, H.-M., Margulies, D. S., Zuo, X.-N., & Holmes, A. J. (2021). Shifting gradients of macroscale cortical organization mark the transition from childhood to adolescence. Proceedings of the National Academy of Sciences, 118(28). doi:10.1073/pnas.2024448118

Donoghue, T., Haller, M., Peterson, E. J., Varma, P., Sebastian, P., Gao, R.,…Voytek, B. (2020). Parameterizing neural power spectra into periodic and aperiodic components. Nat Neurosci, 23(12), 1655–1665. doi:10.1038/s41593-020-00744-x

Duncan, J., Assem, M., & Shashidhara, S. (2020). Integrated Intelligence from Distributed Brain Activity. Trends Cogn Sci, 24(10), 838–852. doi:10.1016/j.tics.2020.06.012

Fedorenko, E., Duncan, J., & Kanwisher, N. (2013). Broad domain generality in focal regions of frontal and parietal cortex. Proc Natl Acad Sci U S A, 110(41), 16616–16621. doi:10.1073/pnas.1315235110

Fox, M. D., Snyder, A. Z., Vincent, J. L., Corbetta, M., Van Essen, D. C., & Raichle, M. E. (2005). The human brain is intrinsically organized into dynamic, anticorrelated functional networks. Proceedings of the National Academy of Sciences, 102(27), 9673–9678. doi:10.1073/pnas.0504136102

Gao, R., Peterson, E. J., & Voytek, B. (2017). Inferring synaptic excitation/inhibition balance from field potentials. Neuroimage, 158, 70–78. doi:10.1016/j.neuroimage.2017.06.078

Glasser, M. F., Coalson, T. S., Robinson, E. C., Hacker, C. D., Harwell, J., Yacoub, E.,…Van Essen, D. C. (2016). A multi-modal parcellation of human cerebral cortex. Nature, 536(7615), 171–178. doi:10.1038/nature18933

Glasser, M. F., Sotiropoulos, S. N., Wilson, J. A., Coalson, T. S., Fischl, B., Andersson, J. L.,…Consortium, W. U.-M. H. (2013). The minimal preprocessing pipelines for the Human Connectome Project. Neuroimage, 80, 105–124. doi:10.1016/j.neuroimage.2013.04.127

Gratton, C., Sun, H., & Petersen, S. E. (2018). Control networks and hubs. Psychophysiology, 55(3). doi:10.1111/psyp.13032

Gyurkovics, M., Clements, G. M., Low, K. A., Fabiani, M., & Gratton, G. (2022). Stimulus-induced changes in 1/f-like background activity in EEG. J Neurosci, 42(37), 7144–7151. doi:10.1523/JNEUROSCI.0414-22.2022

He, B. J. (2014). Scale-free brain activity: past, present, and future. Trends Cogn Sci, 18(9), 480–487. doi:10.1016/j.tics.2014.04.003

Hebart, M. N., & Baker, C. I. (2018). Deconstructing multivariate decoding for the study of brain function. Neuroimage, 180(Pt A), 4–18. doi:10.1016/j.neuroimage.2017.08.005

Huntenburg, J. M., Bazin, P. L., & Margulies, D. S. (2018). Large-Scale Gradients in Human Cortical Organization. Trends Cogn Sci, 22(1), 21–31. doi:10.1016/j.tics.2017.11.002

Jacob, M. S., Roach, B. J., Sargent, K. S., Mathalon, D. H., & Ford, J. M. (2021). Aperiodic measures of neural excitability are associated with anticorrelated hemodynamic networks at rest: A combined EEG-fMRI study. Neuroimage, 245, 118705. doi:10.1016/j.neuroimage.2021.118705

Jensen, O., & Mazaheri, A. (2010). Shaping functional architecture by oscillatory alpha activity: gating by inhibition. Front Hum Neurosci, 4, 186. doi:10.3389/fnhum.2010.00186

Jia, S., Liu, D., Song, W., Beste, C., Colzato, L., & Hommel, B. (2024). Tracing conflict-induced cognitive-control adjustments over time using aperiodic EEG activity. Cereb Cortex, 34(5). doi:10.1093/cercor/bhae185

Kałamała, P., Gyurkovics, M., Bowie, D. C., Clements, G. M., Low, K. A., Dolcos, F.,…Gratton, G. (2024). Event-induced modulation of aperiodic background EEG: Attention-dependent and age-related shifts in E:I balance, and their consequences for behavior. Imaging Neuroscience, 2, 1–18. doi:10.1162/imag_a_00054

Karimi-Rouzbahani, H., Rich, A. N., & Woolgar, A. (2026). Spatiotemporal characterisation of information coding in the multiple demand network. Imaging Neurosci (Camb), 4. doi:10.1162/IMAG.a.1171

Kriegeskorte, N., Mur, M., & Bandettini, P. (2008). Representational similarity analysis - connecting the branches of systems neuroscience. Front Syst Neurosci, 2, 4. doi:10.3389/neuro.06.004.2008

Lu, R. (2025). Linking the multiple-demand cognitive control system to human electrophysiological activity. Neuropsychologia, 210, 109096. doi:10.1016/j.neuropsychologia.2025.109096

Lu, R., Dermody, N., Duncan, J., & Woolgar, A. (2024). Aperiodic and oscillatory systems underpinning human domain-general cognition. Communications Biology, 7, 1643. doi:10.1038/s42003-024-07397-7

Lu, R., Pollitt, E., & Woolgar, A. (2025). Distinct and complementary mechanisms of oscillatory and aperiodic alpha activity in visuospatial attention. Imaging Neuroscience, 3. doi:10.1162/IMAG.a.1038

Manning, J. R., Jacobs, J., Fried, I., & Kahana, M. J. (2009). Broadband shifts in local field potential power spectra are correlated with single-neuron spiking in humans. J Neurosci, 29(43), 13613–13620. doi:10.1523/JNEUROSCI.2041-09.2009

Margulies, D. S., Ghosh, S. S., Goulas, A., Falkiewicz, M., Huntenburg, J. M., Langs, G.,…Smallwood, J. (2016). Situating the default-mode network along a principal gradient of macroscale cortical organization. Proc Natl Acad Sci U S A, 113(44), 12574–12579. doi:10.1073/pnas.1608282113

Markello, R. D., Hansen, J. Y., Liu, Z. Q., Bazinet, V., Shafiei, G., Suarez, L. E.,…Misic, B. (2022). neuromaps: structural and functional interpretation of brain maps. Nat Methods, 19(11), 1472–1479. doi:10.1038/s41592-022-01625-w

Moerel, D., Rich, A. N., & Woolgar, A. (2024). Selective attention and decision-making have separable neural bases in space and time. J Neurosci. doi:10.1523/JNEUROSCI.0224-24.2024

Murphy, A. C., Bertolero, M. A., Papadopoulos, L., Lydon-Staley, D. M., & Bassett, D. S. (2020). Multimodal network dynamics underpinning working memory. Nat Commun, 11(1), 3035. doi:10.1038/s41467-020-15541-0

Nee, D. E. (2021). Integrative frontal-parietal dynamics supporting cognitive control. Elife, 10. doi:10.7554/eLife.57244

Ouyang, G., Hildebrandt, A., Schmitz, F., & Herrmann, C. S. (2020). Decomposing alpha and 1/f brain activities reveals their differential associations with cognitive processing speed. Neuroimage, 205, 116304. doi:10.1016/j.neuroimage.2019.116304

Paquola, C., Bethlehem, R. A., Seidlitz, J., Wagstyl, K., Romero-Garcia, R., Whitaker, K. J.,…Bullmore, E. T. (2019). Shifts in myeloarchitecture characterise adolescent development of cortical gradients. Elife, 8. doi:10.7554/eLife.50482

Pavlov, Y. G., & Kotchoubey, B. (2022). Oscillatory brain activity and maintenance of verbal and visual working memory: A systematic review. Psychophysiology, 59(5), e13735. doi:10.1111/psyp.13735

Pei, L., Northoff, G., & Ouyang, G. (2023). Comparative analysis of multifaceted neural effects associated with varying endogenous cognitive load. Commun Biol, 6(1), 795. doi:10.1038/s42003-023-05168-4

Robinson, E. C., Garcia, K., Glasser, M. F., Chen, Z., Coalson, T. S., Makropoulos, A.,…Rueckert, D. (2018). Multimodal surface matching with higher-order smoothness constraints. Neuroimage, 167, 453–465. doi:10.1016/j.neuroimage.2017.10.037

Robinson, E. C., Jbabdi, S., Glasser, M. F., Andersson, J., Burgess, G. C., Harms, M. P.,…Jenkinson, M. (2014). MSM: a new flexible framework for Multimodal Surface Matching. Neuroimage, 100, 414–426. doi:10.1016/j.neuroimage.2014.05.069

Salimi-Khorshidi, G., Douaud, G., Beckmann, C. F., Glasser, M. F., Griffanti, L., & Smith, S. M. (2014). Automatic denoising of functional MRI data: combining independent component analysis and hierarchical fusion of classifiers. Neuroimage, 90, 449–468. doi:10.1016/j.neuroimage.2013.11.046

Schultz, D. H., Ito, T., & Cole, M. W. (2022). Global Connectivity Fingerprints Predict the Domain Generality of Multiple-Demand Regions. Cereb Cortex. doi:10.1093/cercor/bhab495

Shafiei, G., Baillet, S., & Misic, B. (2022). Human electromagnetic and haemodynamic networks systematically converge in unimodal cortex and diverge in transmodal cortex. PLoS Biol, 20(8), e3001735. doi:10.1371/journal.pbio.3001735

Spreng, R. N., Wojtowicz, M., & Grady, C. L. (2010). Reliable differences in brain activity between young and old adults: a quantitative meta-analysis across multiple cognitive domains. Neurosci Biobehav Rev, 34(8), 1178–1194. doi:10.1016/j.neubiorev.2010.01.009

Sternberg, S. (1966). High-Speed Scanning in Human Memory. Science, 153(3736), 652–654. doi:10.1126/science.153.3736.652

Stigler, J. W., Lee, S.-Y., & Stevenson, H. W. (1986). Digit memory in Chinese and English: Evidence for a temporally limited store. Cognition, 23(1), 1–20. doi:10.1016/0010-0277(86)90051-X

Suarez, L. E., Markello, R. D., Betzel, R. F., & Misic, B. (2020). Linking Structure and Function in Macroscale Brain Networks. Trends Cogn Sci, 24(4), 302–315. doi:10.1016/j.tics.2020.01.008

Sydnor, V. J., Larsen, B., Bassett, D. S., Alexander-Bloch, A., Fair, D. A., Liston, C.,…Satterthwaite, T. D. (2021). Neurodevelopment of the association cortices: Patterns, mechanisms, and implications for psychopathology. Neuron. doi:10.1016/j.neuron.2021.06.016

Thuwal, K., Banerjee, A., & Roy, D. (2021). Aperiodic and Periodic Components of Ongoing Oscillatory Brain Dynamics Link Distinct Functional Aspects of Cognition across Adult Lifespan. eNeuro, 8(5). doi:10.1523/ENEURO.0224-21.2021

Ugurbil, K., Xu, J., Auerbach, E. J., Moeller, S., Vu, A. T., Duarte-Carvajalino, J. M.,…Consortium, W. U.-M. H. (2013). Pushing spatial and temporal resolution for functional and diffusion MRI in the Human Connectome Project. Neuroimage, 80, 80–104. doi:10.1016/j.neuroimage.2013.05.012

van Engen, Q., Chau, G., Smith, A., Adam, K., Donoghue, T., & Voytek, B. (2026). Dissociating Contributions of Theta and Alpha Oscillations from Aperiodic Neural Activity in Human Visual Working Memory. J Neurosci, 46(7). doi:10.1523/JNEUROSCI.2340-24.2025

Vasa, F., & Misic, B. (2022). Null models in network neuroscience. Nat Rev Neurosci, 23(8), 493–504. doi:10.1038/s41583-022-00601-9

Vazquez-Rodriguez, B., Suarez, L. E., Markello, R. D., Shafiei, G., Paquola, C., Hagmann, P.,…Misic, B. (2019). Gradients of structure-function tethering across neocortex. Proc Natl Acad Sci U S A, 116(42), 21219–21227. doi:10.1073/pnas.1903403116

Vinck, M., Uran, C., Dowdall, J. R., Rummell, B., & Canales-Johnson, A. (2024). Large-scale interactions in predictive processing: oscillatory versus transient dynamics. Trends Cogn Sci. doi:10.1016/j.tics.2024.09.013

Voytek, B., & Knight, R. T. (2015). Dynamic network communication as a unifying neural basis for cognition, development, aging, and disease. Biol Psychiatry, 77(12), 1089–1097. doi:10.1016/j.biopsych.2015.04.016

Waschke, L., Donoghue, T., Fiedler, L., Smith, S., Garrett, D. D., Voytek, B., & Obleser, J. (2021). Modality-specific tracking of attention and sensory statistics in the human electrophysiological spectral exponent. Elife, 10, e70068. doi:10.7554/eLife.70068

Wen, H., & Liu, Z. (2016a). Broadband Electrophysiological Dynamics Contribute to Global Resting-State fMRI Signal. J Neurosci, 36(22), 6030–6040. doi:10.1523/JNEUROSCI.0187-16.2016

Wen, H., & Liu, Z. (2016b). Separating Fractal and Oscillatory Components in the Power Spectrum of Neurophysiological Signal. Brain Topogr, 29(1), 13–26. doi:10.1007/s10548-015-0448-0

Woolrich, M. W., Ripley, B. D., Brady, M., & Smith, S. M. (2001). Temporal autocorrelation in univariate linear modeling of FMRI data. Neuroimage, 14(6), 1370–1386. doi:10.1006/nimg.2001.0931

Yan, J., Yu, S., Muckschel, M., Colzato, L., Hommel, B., & Beste, C. (2024). Aperiodic neural activity reflects metacontrol in task-switching. Sci Rep, 14(1), 24088. doi:10.1038/s41598-024-74867-7

Yeo, B. T., Krienen, F. M., Sepulcre, J., Sabuncu, M. R., Lashkari, D., Hollinshead, M.,…Buckner, R. L. (2011). The organization of the human cerebral cortex estimated by intrinsic functional connectivity. J Neurophysiol, 106(3), 1125–1165. doi:10.1152/jn.00338.2011

Zheng, Y., Lu, R., & Woolgar, A. (2024). Radical flexibility of neural representation in frontoparietal cortex and the challenge of linking it to behaviour. Current Opinion in Behavioral Sciences, 57, 101392. doi:10.1016/j.cobeha.2024.101392

